# SARS-CoV-2 infection of human iPSC-derived cardiac cells predicts novel cytopathic features in hearts of COVID-19 patients

**DOI:** 10.1101/2020.08.25.265561

**Authors:** Juan A. Pérez-Bermejo, Serah Kang, Sarah J. Rockwood, Camille R. Simoneau, David A. Joy, Gokul N. Ramadoss, Ana C. Silva, Will R. Flanigan, Huihui Li, Ken Nakamura, Jeffrey D. Whitman, Melanie Ott, Bruce R. Conklin, Todd C. McDevitt

**Affiliations:** Gladstone Institutes, San Francisco, CA; Biomedical Sciences PhD Program, University of California, San Francisco, CA; UC Berkeley UCSF Joint Program in Bioengineering, Berkeley, CA; UCSF Department of Neurology, San Francisco, CA; UCSF Department of Laboratory Medicine, San Francisco, CA; Innovative Genomics Institute, Berkeley, CA; UCSF Department of Ophthalmology, San Francisco, CA; UCSF Department of Medicine, San Francisco, CA; UCSF Department of Bioengineering and Therapeutic Sciences, San Francisco, CA

## Abstract

Although COVID-19 causes cardiac dysfunction in up to 25% of patients, its pathogenesis remains unclear. Exposure of human iPSC-derived heart cells to SARS-CoV-2 revealed productive infection and robust transcriptomic and morphological signatures of damage, particularly in cardiomyocytes. Transcriptomic disruption of structural proteins corroborated adverse morphologic features, which included a distinct pattern of myofibrillar fragmentation and numerous iPSC-cardiomyocytes lacking nuclear DNA. Human autopsy specimens from COVID-19 patients displayed similar sarcomeric disruption, as well as cardiomyocytes without DNA staining. These striking cytopathic features provide new insights into SARS-CoV-2 induced cardiac damage, offer a platform for discovery of potential therapeutics, and raise serious concerns about the long-term consequences of COVID-19.

## INTRODUCTION

Initial descriptions of COVID-19, the pandemic disease caused by severe acute respiratory syndrome coronavirus 2 (SARS-CoV-2), characterized it as a primarily respiratory syndrome^1^. However, increasing clinical evidence now implicates multiple organ systems in COVID-19, including the heart, gastrointestinal tract, and kidneys^2–5^. Notably, multiple independent reports have reported cases of acute COVID-19 associated myopathy^6–8^, even without prior cardiovascular disease^9^, indicating that SARS-CoV-2 may be directly causing cardiac damage. Meta-analyses identify elevated high-sensitivity troponin-I or natriuretic peptides, biomarkers of cardiac damage, as the strongest predictors of mortality in hospitalized patients, eclipsing both prior congestive obstructive pulmonary disease and cardiovascular disease^8–11^. Alarmingly, 55% of COVID-19 patients presented abnormal echocardiograms^12^, and a majority of recovered patients continue to suffer from impaired cardiac function, indicating that long-term heart sequelae from COVID-19 may not be limited to severe cases^13^.

Identifying therapeutic strategies to prevent or manage myocardial injury in COVID-19 patients is impeded by a limited understanding of the mechanisms by which SARS-CoV-2 induces cardiac damage. Cardiac damage may be caused by systemic impacts of SARS-CoV-2, such as hypoxic stress due to pulmonary damage, microvascular thrombosis, and/or the systemic immune response to viral infection^14^. However, cardiomyocytes are known to express the primary receptor for viral entry, ACE2^15^ and thus may be infectable by SARS-CoV-2^16^ Viral RNA has been detected in myocardial autopsies of SARS-CoV^17^ and SARS-CoV-2^18^ infected patients, and viral particles have been found inside cardiomyocytes and other cardiac cells in COVID-19 patients^19,20,21^, suggesting that direct myocardial infection may cause COVID-19 cardiac injury.

Despite the alarming clinical consequences of COVID-19 in the heart, pathological studies of patient autopsy samples have not described any specific effect in myocardial specimens, apart from diffuse edema and occasional hypertrophy^22–24^. Detailed pathological studies have been hampered by biosafety considerations and the limitations of hematoxylin and eosin (H&E). staining. In addition, sample availability is restricted to post-mortem specimens, which limits most observations to late stage disease endpoints.

*Ex vivo* studies using human cell-based models of the heart, such as cardiac tissue derived from human induced pluripotent stem cells (iPSCs), afford the most direct route for prospective and clinically-relevant studies on the effects of cardiac viral infection. Stem-cell derived models have already demonstrated the susceptibility of hepatocytes^25^, intestinal epithelium^26,27^, and lung organoids^28^ to SARS-CoV-2 infection. While two recent reports confirmed that human iPSC-cardiomyocytes are susceptible to SARS-CoV-2 infection^29,30^, specific cardiac cytopathic features have yet to be identified. In addition, the viral tropism for other cardiac cell types, which may be involved in microthrombosis^31^ or weakening of the ventricular wall, has not been explored, nor has there been direct correlation of *in vitro* results to clinical pathology specimens. Here, we examined the relative susceptibility of three cardiac cell types derived from iPSCs: cardiomyocytes (CMs), cardiac fibroblasts (CFs), and endothelial cells (ECs), to SARS-CoV-2 infection, and identified clear hallmarks of infection and cardiac cytopathy that predict pathologic features found in human COVID-19 tissue specimens. The observed phenotypic biomarkers of SARS-CoV-2 infection in cardiomyocytes and associated discovery platform can enable rapid development of cardioprotective therapies for COVID-19.

## RESULTS

### SARS-CoV-2 infects and propagates in cardiomyocytes, but not endothelial cells or cardiac fibroblasts

The relative susceptibility of different cardiac cell types to SARS-CoV-2 infection has not been characterized, leading to ambiguity over the sources of cardiac damage and relevant therapeutic targets. Analysis of single-cell RNA-sequencing and immunofluorescence staining of ECs, CFs, and CMs revealed that ACE2, the receptor for SARS-CoV-2, was only detectable in CMs **(Supplementary Information 1, Supp. Figs. 1A-D)**. Although the cell surface protease TMPRSS2, commonly involved in viral cell entry, was not detected in any cell type, cathepsin-L (CTSL) and cathepsin-B (CTSB) were detected in all cells. These observations suggest potential infectivity of CMs by SARS-CoV-2, and predict poor infectivity in ECs and CFs. To validate these predictions, we exposed human iPSC-derived CMs, CFs, ECs, or a mixture of all three to mimic native myocardial composition, to SARS-CoV-2 at a low MOI (MOI = 0.006).

**Figure 1.**
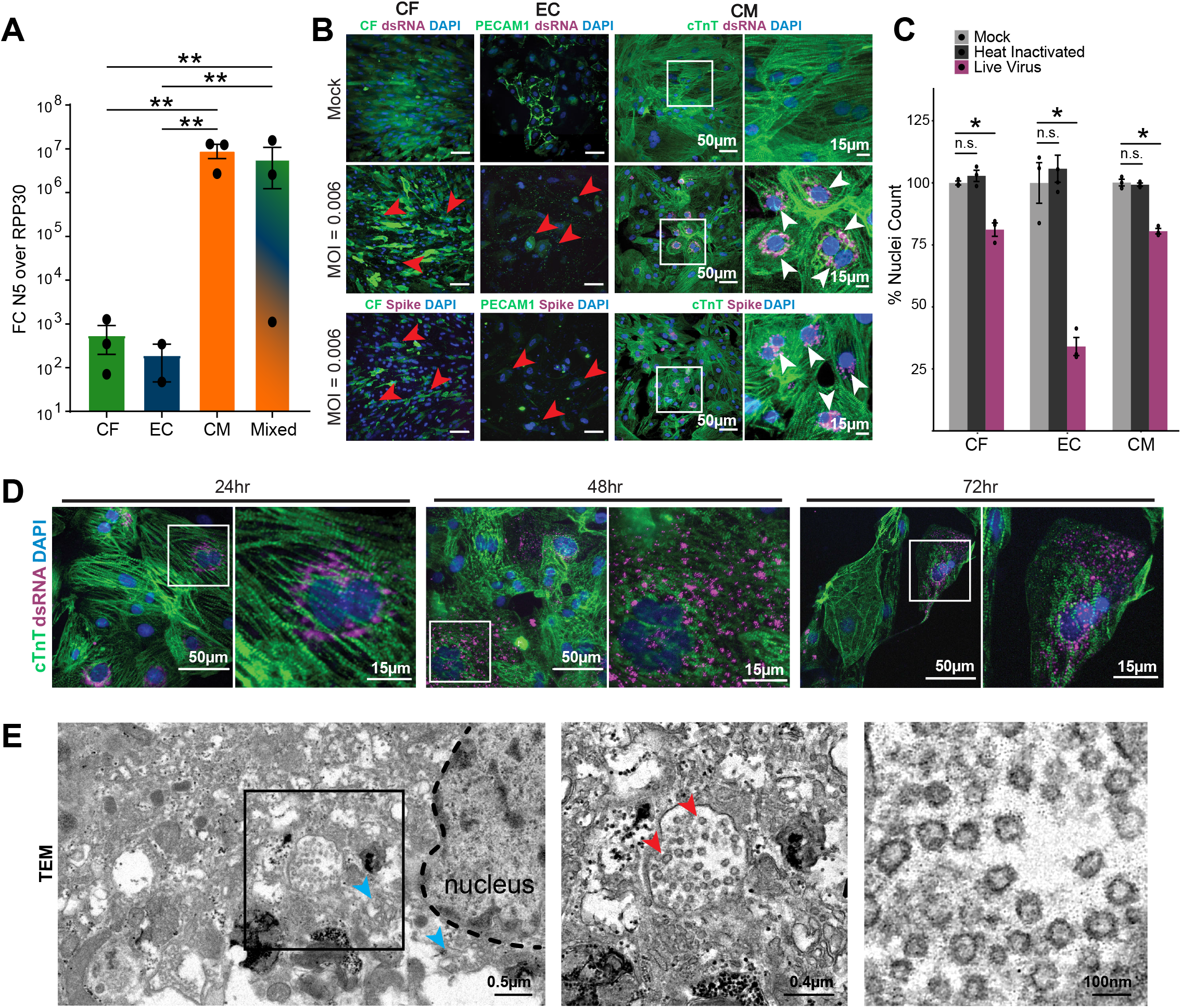
Effects of SARS-CoV-2 exposure on different iPS-derived cardiac cell types. In all experiments, cells were exposed to SARS-CoV-2 virus for 48 hours, unless otherwise specified, at an MOI of 0.006 before lysis or fixation. **A**. RT-qPCR quantification of viral RNA (Fold change of SARS-CoV-2 N gene, N5, over housekeeping gene transcript, RPP30) in cell cultures exposed to SARS-CoV-2. CF: iPSC-derived cardiac fibroblasts; EC: iPSC-derived endothelial cells; CM: iPSC-derived cardiomyocytes; Mixed: 60:30:10 CM:EC:CF. Error bars: SEM. **: p-val < 0.01, one-way ANOVA with Tukey’s multiple comparisons. **B**. Representative images of immunostaining of cardiac cells exposed to SARS-CoV-2. PECAM-1 (CD31) was used as an endothelial cell marker, and cTnT as a cardiomyocyte specific marker. Cardiac fibroblast (CF) cells expressed GFP constitutively. Viral signal was detected by staining for SARS-CoV-2 Spike protein or viral double stranded RNA (dsRNA), as noted. White boxes represent zoomed in regions in rightmost panels. White arrows denote clusters of dsRNA signal. **C**. Toxicity of SARS-CoV-2 to cardiac cell types, quantified by nuclear retention. Y-axis depicts the % of nuclei counted (relative to mock). Nuclei were counted automatically at 10x magnification (10 images/condition). Light gray: Vehicle treatment (mock), Dark gray: Heat inactivated SARS-CoV-2 (MOI = 0.1), Magenta: SARS-CoV-2 (MOI = 0.006). **D**. Representative images of immunostaining of infected cardiomyocytes at 24h, 48h or 72h after addition of SARS-CoV-2 virus. White boxes indicate zoomed in regions. **E**. Transmission electron microscopy of SARS-CoV-2 viral particles in an infected CM. Left: Montage view with nucleus (dashed line), in addition to remnant ER-Golgi (light blue arrowheads), with viral particles enclosed in a membrane compartment. Middle: Close up of SARS-CoV-2 virions (red arrowheads) and surrounding membrane. Right: Magnified view of SARS-CoV-2 virions, showing the 500-750nm diameter membrane and the 60-100nm diameter viral particles within.

After 48 hours, CFs and ECs showed little to no viral RNA relative to a housekeeping control, whereas CMs expressed >10^4^ greater levels of viral RNA than CFs or ECs **(Fig. 1A)**. There was no significant difference in viral detection between CMs and mixed cultures **(Fig. 1A)**, and undifferentiated iPSCs were uninfectable **(Supp. Fig. S1D)**. Differences in viral expression largely correlated with cell-type specific ACE2 expression levels **(Supp. Figs. 1A-B)**. Plaque assays on the supernatants of virally exposed cells confirmed that CFs, ECs, and iPSCs did not support productive infection, whereas CMs robustly produced replication-competent virions **(Supp. Fig. S1E)**.

Immunostaining for viral Spike protein and for double-stranded RNA (dsRNA), indicative of replicating virus, further confirmed that only CMs supported viral replication **(Fig. 1B, white arrows)**. However, all three cell types exhibited significant cytopathic effects after 48 hours of viral exposure, characterized by fragmented cell bodies, dissociation from neighboring cells **(Fig. 1B, red arrows)** and significant cell death **(Fig. 1C)**. Visual cytopathic effects were most prevalent in CFs, while the greatest nuclear loss was observed in ECs, indicating toxicity from viral exposure can occur without detectable viral replication. However, inoculation with heat-inactivated SARS-CoV-2 did not cause cell death or dissociation in any of the cell types assayed **(Fig. 1C)**.

Replication of positive-strand single stranded RNA viruses, including SARS-CoV-2, involves budding of double-membrane vesicles from the endoplasmic reticulum, with viral particle assembly occurring in cisternae of the ER-Golgi intermediate compartment (ERGIC)^32^. In CMs infected with SARS-CoV-2, dsRNA and Spike protein initially (24h post-infection) accumulated near the nucleus in small perinuclear puncta, the typical location of the ERGIC, indicating potential replication centers **(Fig. 1D)**. At 48h post infection, many cells exhibited dsRNA signal dispersed throughout their cytoplasm, which may correlate with advanced stages of infection. By 72h post infection, SARS-CoV-2 was spread throughout the culture and a large portion of the CMs had died, with the remaining cells displaying disperse viral stain localization, dissociation from neighboring cells, and heavily reduced sarcomeric staining **(Fig. 1D)**.

Remnants of the ER-Golgi membranes (light blue arrowheads) and large vesicles near the nucleus in infected CMs were readily identified by transmission electron microscopy (TEM). These vesicles, approximately 500-750 nanometers in diameter, contained multiple particles 60-100 nm in diameter that were identified as SARS-CoV-2 virions **(Fig. 1E, red arrowheads)**. Altogether, these results indicate that SARS-CoV-2 is able to infect, replicate in, and rapidly propagate through CMs.

### SARS-CoV-2 infection of cardiomyocytes is dependent on an endolysosomal route

We next sought to elucidate the mechanism of SARS-CoV-2 entry into CMs. Pretreatment of cells with an ACE2 blocking antibody or with cathepsin-B and -L inhibitor E64d significantly reduced viral detection in infected CMs **(Fig. 2A, S1F)**. Though FURIN is expressed in CMs **(Fig. S1A)**, FURIN inhibition did not reduce infection **(Fig. S1F)**. Inhibiting specific cathepsins revealed that CTSL inhibition via Z-Phe-Tyr(tBu)-diazomethylketone (Z-FY-DK) decreased viral detection in infected cells to approximately 10% of vehicle-treated controls, whereas inhibition of CTSB with CA-074 did not attenuate viral levels **(Fig. 2B)**. In addition, the PIKfyve inhibitor apilimod and autolysosome acidification blocker bafilomycin each successfully reduced viral infection to ∼0.1% or 1% of vehicle-treated controls, respectively. In contrast, the TMPRSS2 inhibitors aprotinin and camostat mesilate did not significantly inhibit viral infection. These results suggest that SARS-CoV-2 binds to iPS-CMs via the ACE2 receptor, and utilizes a CTSL (but not CTSB)-dependent endolysosomal route to infection, independent of TMPRSS2/serine protease-mediated activation at the cellular membrane.

**Figure 2.**
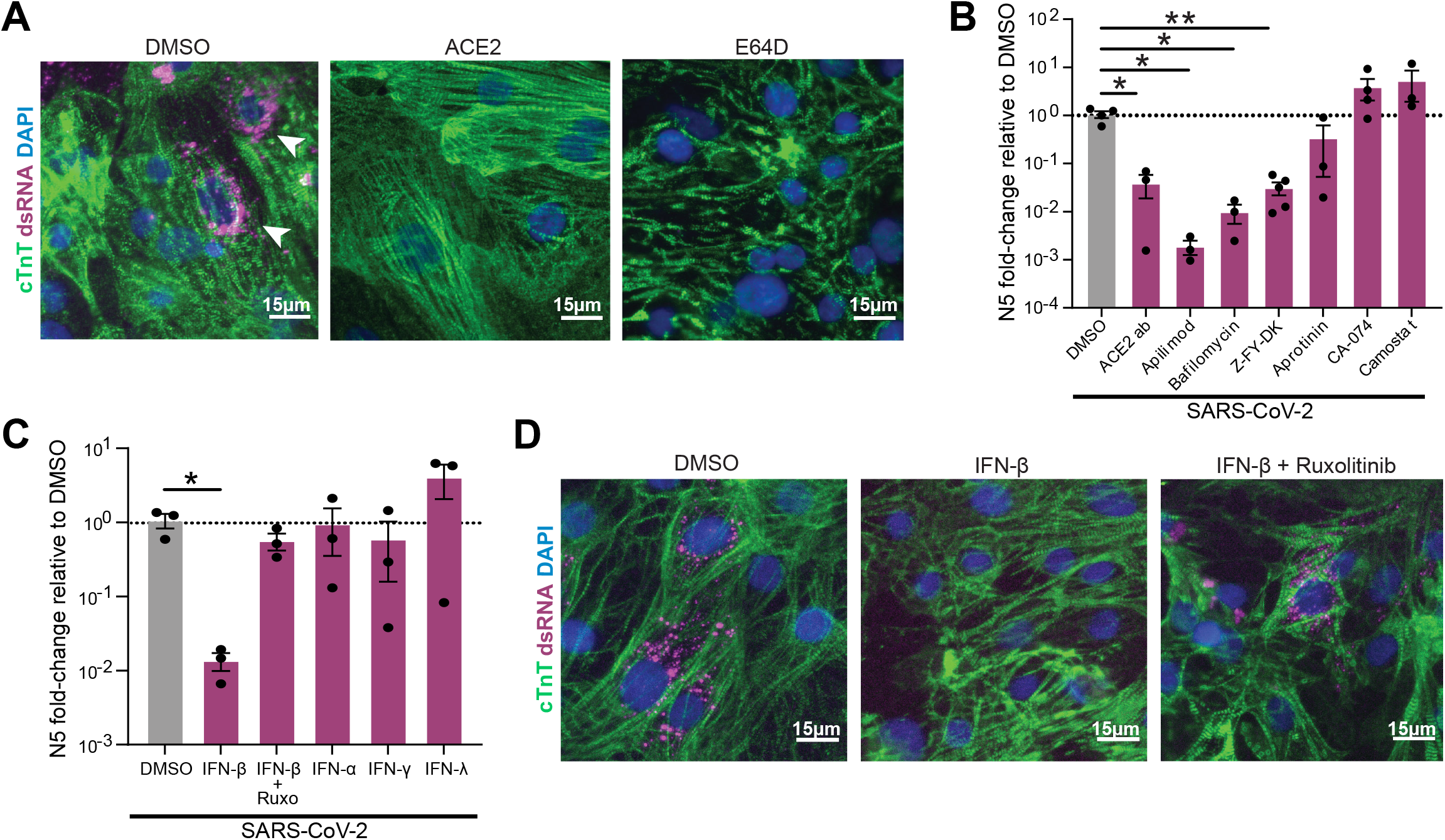
Pharmacological modulation of SARS-CoV-2 infection and innate immune response in CMs. **A**. Representative immunofluorescence images from SARS-CoV-2 infected (MOI=0.006) CMs pretreated with either vehicle (DMSO), ACE2 blocking antibody (‘ACE2ab’) or cathepsin-B and -L inhibitor E64d for 2h before infection. Double-stranded RNA (dsRNA) staining (white arrowheads) denotes presence of replicating virus. **B-C**. RT-qPCR quantification of viral RNA (N5) in CM samples exposed to SARS-CoV-2 for 48h (MOI=0.006) after 2h pretreatment with the indicated reagents to block viral entry (**B**) or prime the cells’ innate immune response (**C**). Dots represent separate replicates. *: p-val < 0.05, **: p-val < 0.01. *N*>=3 for all conditions. One-way ANOVA with Tukey’s multiple comparisons. Z-FY-DK: Z-Phe-Tyr(tBu)-diazomethylketone, specific Cathepsin-L inhibitor; CA-074: specific Cathepsin-B inhibitor; Ruxo: Ruxolitinib, JAK1/2 inhibitor. **D**. Representative immunofluorescence images from cardiomyocytes pre-treated with vehicle (DMSO), IFN-β, or IFN-β with JAK inhibitor ruxolitinib.

We next examined whether priming the innate immune response could reduce SARS-CoV-2 infection of CMs. CMs were treated with IFN-α, IFN-β, IFN-γ, or IFN-λ prior to infection. Only pre-exposure to IFN-β decreased infection, and this effect was reversed by co-treatment with the JAK/Stat inhibitor ruxolitinib **(Figs. 2C-D)**. Our observation that IFN-β but not IFN-α pretreatment reduced infection is in concordance with a previous report of differential antiviral activity of these two type-I interferons in a mouse model of myocarditis^33^, confirming that CMs can mount an antiviral response with appropriate stimulation.

### SARS-CoV-2 exposure induces transcriptional changes in genes involved in contractile machinery

To evaluate the transcriptional impact of SARS-CoV-2 on cardiac cells, we performed RNA-sequencing of CFs, ECs, and iPSCs exposed to an MOI of 0.006 and CMs exposed to a range of elevating MOIs (0.001, 0.01, and 0.1). Sequencing recovered SARS-CoV-2 reads in an MOI- and cell type-dependent fashion **(Fig. 3A)**, with SARS-CoV-2 reaching >50% of the recovered reads in CMs at the highest MOI. Principal component analysis (PCA) of the biological conditions revealed clustering based primarily on cell type, with CFs and ECs clustering together and CMs and iPSCs separating into distinct clusters **(Fig. 3B)**. Loading plots of the principal components supported this interpretation: genes determining the spectrum of variation between CMs and CF/ECs were associated with CMs (*MYH7, MYH6, TNNT2*) at one pole **(Fig. 3C)** and anti-correlated with CF/EC specific genes at the other (*FN1, COL1A2, TFPI2, MME*) **(Supplementary Fig. S2A)**. However, the distance between mock-treated CMs and the furthest infected CMs was greater than the distance between CMs and CFs or ECs, indicating that viral infection altered cellular expression profiles at least as strongly as cell type. Along this axis, the level of transcriptional disruption correlated poorly with MOI across all CM samples, potentially due to natural stochasticity in the kinetics of infection. Regrouping conditions by the level of transcriptional disruption allowed us to more clearly deduce transcriptional trends resulting from viral exposure **(Figs. S2A-D)**. Infected cardiomyocytes downregulated pathways corresponding to cardiac muscle tissue organization and cellular respiration. Meanwhile, pathways related to the innate immune response and apoptosis were upregulated, although the innate response to viral infection decreased, suggesting active repression by SARS-CoV-2 as reported by others^34^**(Fig. 3D)**. Anomalous upregulation of pathways associated with olfactory receptors and dysregulation of proteasome catabolism were also observed with increasing transcriptional disruption **(Fig. 3D, S2E)**.

**Figure 3.**
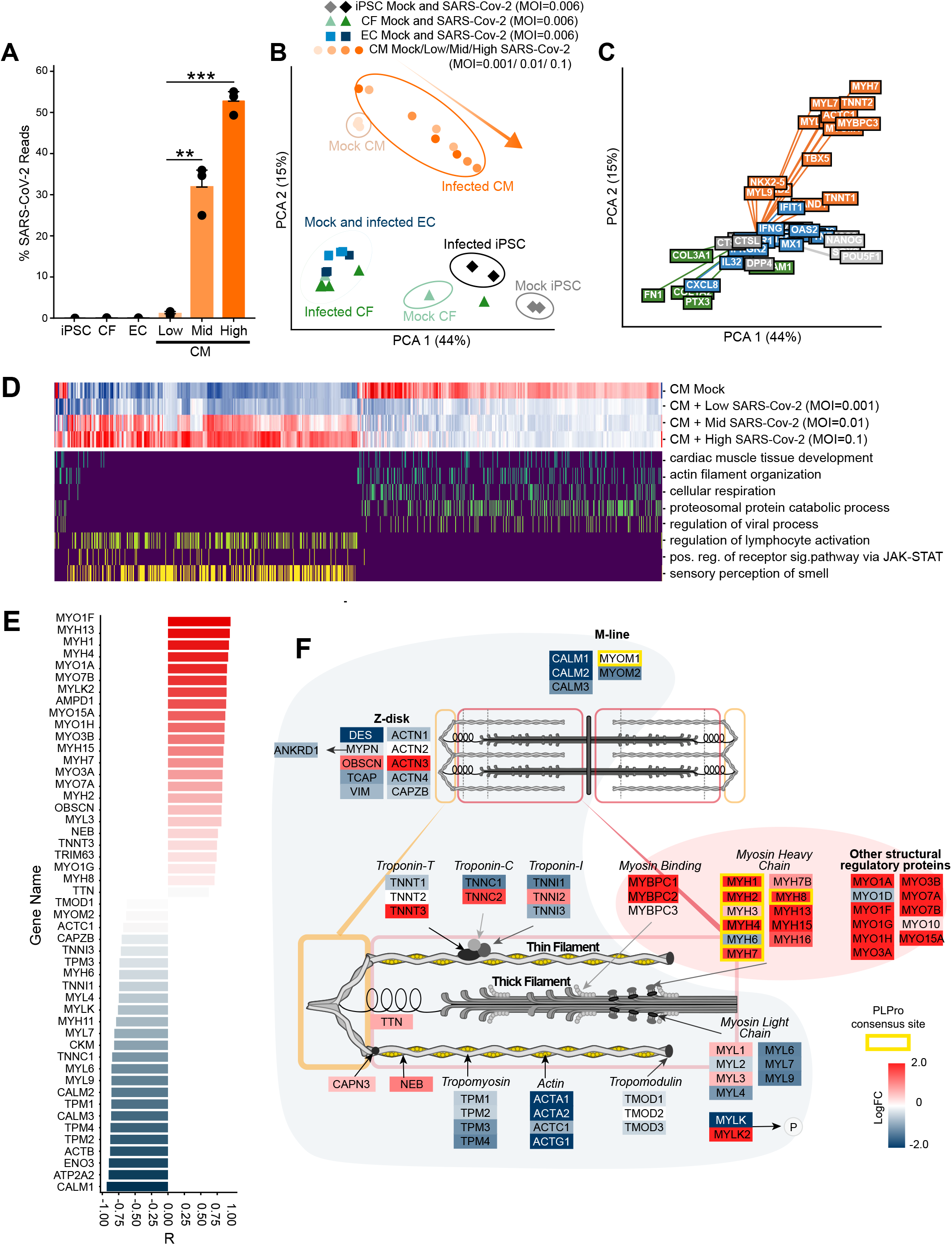
Transcriptional effects of SARS-CoV-2 exposure to cardiac cells. **A**. Percentage of total reads that map to the SARS-CoV-2 viral genome in various cell types. iPSCs, ECs or CFs were exposed at an MOI of 0.006, and CMs at three different MOIs: 0.001 (‘Low’), 0.01 (‘Mid’) and 0.1 (‘High’). **: p-val < 0.01; ***: p-val < 0.001. **B**. Principal component analysis of transcriptomic samples. Dot shapes and colors represent the different cell types, whether they were exposed to SARS-CoV-2 virus and, in the case of CMs, the different MOIs used. **C**. Loading plot for a selected subset of genes, with color indicating cardiomyocyte state (orange), fibroblast/endothelial cell state (green), iPSC state (light gray), SARS-CoV-2 infection-related factors (dark gray), immune response (blue). **D**. Heat map depicting transcriptional expression profiles for mock-infected CMs compared to the least (Low Inf), middle (Mid Inf), and most (High Inf) transcriptionally disrupted CM samples for genes mapping to GO terms of interest (genes |log2 fold change| > 1 between high infection and mock, FDR < 0.05; GO terms containing at least 25 enriched genes, FDR < 0.01). **E**. Expression ratio of genes involved in sarcomeric structure and myosin contractility of the high-infection CM groups relative to the mock-infection CM group. **F**. Schematic drawing of a sarcomere showing localization of differentially regulated factors. Yellow outlines denote proteins which possess the putative SARS-CoV papain-like protease cleavage site LKGGK. Background colors denote log2FC of the high-infection condition over mock. Myosin heavy chains and unconventional myosins are upregulated but almost all thin filament members and myosin light chains are downregulated.

The differentially regulated genes involved in inflammation and innate immunity reflected the observed preferential infectivity of CMs. Exposed CFs and ECs had a depressed cytokine response compared to CMs at all three MOIs **(Supplementary Fig. S2E)**, while infected CMs were enriched in genes involved in cytokine production and T-cell activation (OAS2, MX1, IFIT1, IL1B, IL6, TNF) **(Supplementary Fig. S2E)**. Interestingly, CMs at each MOI showed clear dysregulation of contractile machinery, proteasomal subunits and ubiquitination (**Supplementary Fig. 3A-G)**. Genes involved in the Linker of Nucleoskeleton and Cytoskeleton (LINC) complex were disrupted, especially calmin and members of the nesprin family, both critical for anchoring the nucleus to the actin cytoskeleton **(Fig S4)**. Furthermore, sarcomeric structural proteins, myosin light chains, and proteasome kinases and chaperones were strongly downregulated, while most myosin heavy chains were significantly upregulated **(Fig. 3E, F, Supplementary Fig. 3A-G)**, suggesting a split effect of SARS-CoV-2 infection on the contractile and structural integrity of CMs.

### SARS-CoV-2 infection disrupts multiple intracellular features of cardiomyocytes

Motivated by the expression changes in structural and contractile genes in our transcriptomic data, we performed high-content imaging of CMs following SARS-CoV-2 infection. We immediately noticed several abnormal structural features in infected CMs that were not seen in mock or heat-inactivated virus-treated CMs. Most notably, we observed widespread myofibrillar disruption throughout the cytoplasm: a unique pattern of specific, periodic cleavage of myofibrils into individual sarcomeric units of identical size without any linear alignment **(Figs. 4A-C, Supplementary Figs. 4A-D)**. Myofibrillar fragmentation was evident as early as 24 hours after infection, and significantly increased after 48 hours of viral exposure, suggesting that fragmentation increases over the course of infection **(Fig. 4B)**. This pattern of myofibrillar fragmentation was present in cells independently of actively replicating virus (as per dsRNA staining; Chi-square test for independence p-value = 0.81) **(Fig. 4C)**.

**Figure 4.**
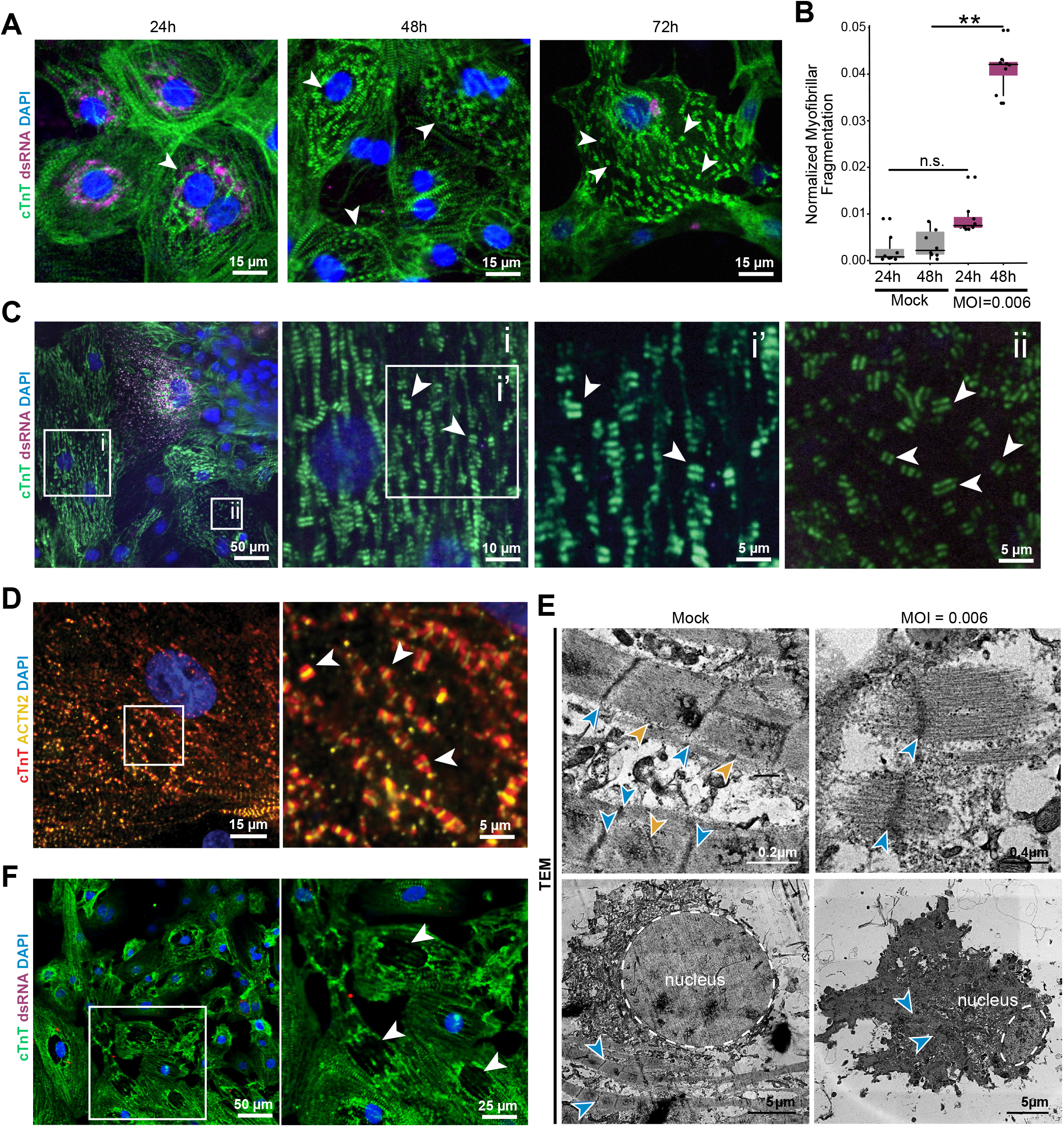
Analysis of cytopathological features induced by SARS-CoV-2 infection in CMs. **A**. Representative immunofluorescence images of myofibrillar fragmentation in CMs at different timepoints after exposure to SARS-CoV-2. White arrowheads indicate fragments consisting of two bands of cTnT positive staining. **B**. Quantification of number of cells presenting myofibrillar fragmentation at 24h and 48h post-exposure (defined as cells presenting at least one event of a cTnT doublet unaligned and dissociated from other myofibrils). Number of cells was normalized to total number of nuclei in the images counted. Each dot represents a separate infection sample. Each replicate is the additive count of 9 randomly acquired fields of view. **: p-val < 0.01. **C**. Representative immunostaining showing a cell staining positively for viral dsRNA, adjacent to cells with different degrees of myofibrillar fragmentation. White squares indicate zoomed-in areas, with labels corresponding to insets. White arrowheads point to examples of cTnT doublets (myofibrillar fragments). **D**. cTnT and ACTN2 double-staining of CMs displaying myofibrillar fragmentation. White arrowheads indicate cTnT-ACTN2-cTnT myofibrillar fragments. **E**. TEM images of sarcomeres in mock treated and SARS-CoV-2 infected (MOI=0.006) CM cultures. Blue arrows denote the sarcomeric z-disk; Yellow arrow indicates M-line location; dashed line delimits nucleus. Sarcomeres of mock treated cells display clear I and A-bands, but fragmented sarcomeres only possess thin filaments. Below: Representative TEM image of a healthy nucleus, and the nucleus of a cell infected with SARS-CoV-2. **F**. Immunofluorescence staining of SARS-CoV-2-exposed CMs displaying loss of nuclear DNA staining (48h post exposure). White arrowheads indicate locations of sarcomeric retraction and absence of nuclear material.

Co-staining of CMs with the thin filament marker cardiac Troponin T (cTnT), and the Z-disk marker α-actinin 2, revealed that SARS-CoV-2-induced myofibrillar fragments consisted of two bands of cTnT flanking a single α-actinin 2 band **(Fig. 4D, Supplementary Figure S5E)**. To examine sarcomeric fragmentation in greater detail, we employed TEM imaging of SARS-CoV-2 infected and mock-treated CMs. While intact sarcomeres were clearly identified with a classic dark Z-disk adjacent to a light I-band followed by a dark A-band, single fragmented myofibrils displayed an extended I-band and complete absence of the A-band **(Fig. 4E)**, indicating a liberation of thick filaments from sarcomere subunits.

Since transcriptomic profiling indicated that viral infection perturbed the proteasome system **(Fig. 3F, Supplementary Fig. S3F)**, we also examined if proteasome inhibition could recapitulate similar structural abnormalities. Although high doses of the proteasome inhibitor bortezomib induced myofibril fragmentations in CMs, the effect was much less frequent and less severe than the effect of SARS-CoV-2 infection, and was generally accompanied by diffuse cTnT staining **(Supplementary Fig. S5G)**. Furthermore, the well-known cardiotoxic drug doxorubicin did not induce myofibril fragmentation (data not shown), suggesting that proteasomal inhibition may specifically recapitulate part of the viral effects that lead to myofibrillar fragmentation. Additionally, we also observed a second structural phenotype where infected CMs often lacked nuclear DNA staining. This phenomenon was characterized by withdrawn sarcomeres and abnormally shaped or absent nuclear DNA signal **(Fig. 4F)**. Interestingly, this abnormal nuclear structure is consistent with our observed transcriptomic dysregulation of the LINC complex **(Supplementary Fig. S4)**.

### *In vitro* findings predict disruptions in myocardium of COVID-19 patients

We next asked whether the SARS-CoV-2 induced phenotypes observed *in vitro* could predict similar patterns of cardiac cell damage *in vivo*. We obtained autopsy specimens from three COVID-19 positive patients: one who had been diagnosed with myocarditis (COVID-M1), and two which had no reported cardiac involvement (COVID-A1 and COVID-A2). The myocardium of COVID-M1 revealed the expected immune infiltration of the tissue, in addition to significant disruptions in myocytes lacking DNA staining **(Fig. 5A-B, red arrowheads; Supplementary Figs. S6A-B)**. Moreover, COVID-A1 **(Fig. 5C)** and COVID-A2 patients **(Supplementary Fig. 6B)** both had loss of nuclear DNA comparable to that of the COVID-M1 patient. A slight difference (approximately 10%) in nuclear counts between regions of intact tissue vs disrupted tissue was detected across patients (**Supplementary Fig. 6C**).

**Figure 5.**
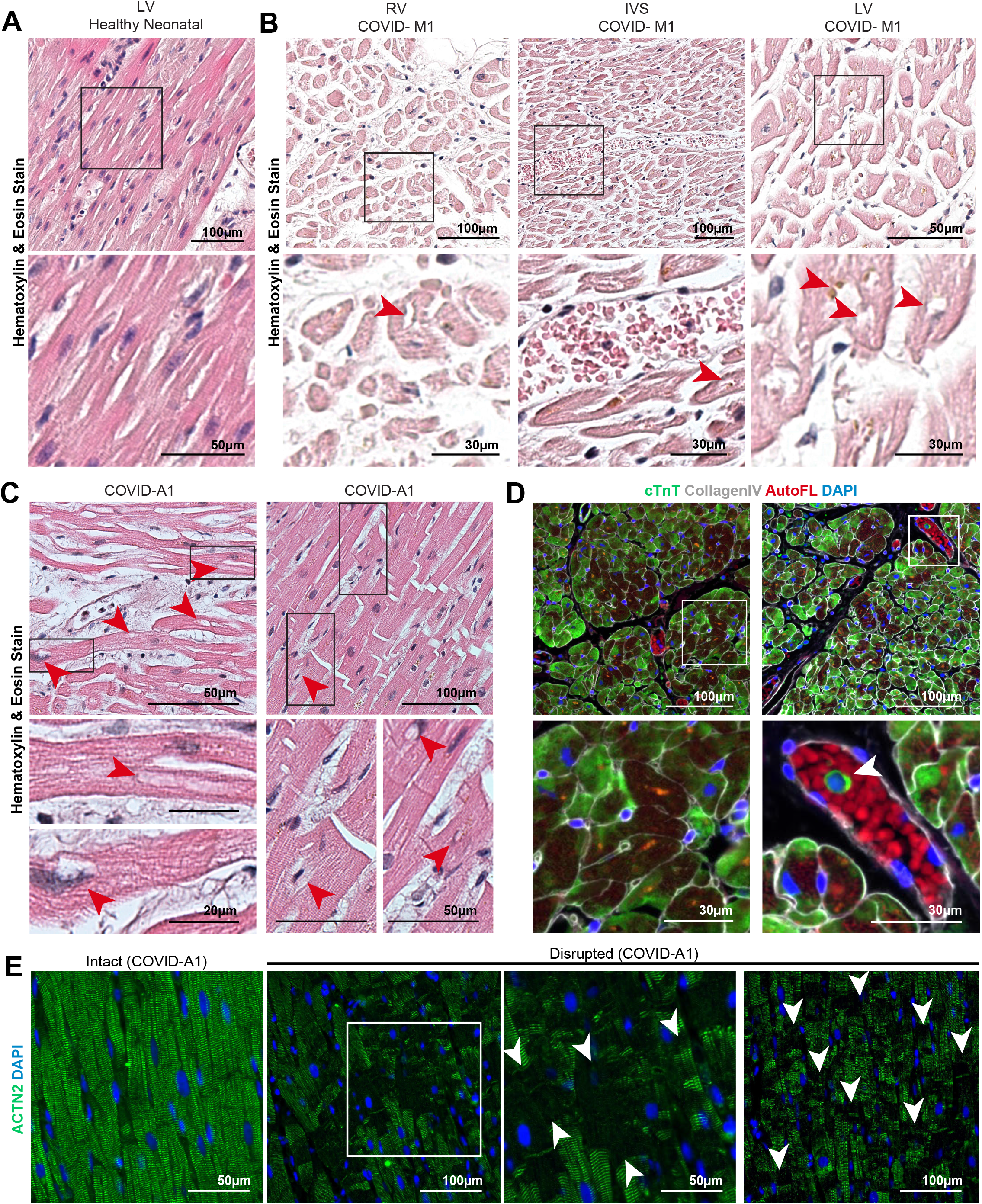
Pathological features of autopsy myocardial tissue from SARS-CoV-2-infected patients. **A**. Hematoxylin and Eosin (H&E) staining of healthy neonatal left ventricle tissue. **B**. H&E staining images of myocardial tissue from a COVID-19 patient with diagnosed myocarditis (COVID-M1). Black boxes indicate zoomed in areas of the bottom panels. Red arrowheads indicate cardiomyocytes lacking chromatin stain. **C**. Representative H&E staining images of myocardial tissue from a COVID-19 patient without diagnosed myocarditis (COVID-A1). Red arrowheads denote putative nuclear locations lacking chromatin stain. **D** Representative immunofluorescence staining of the myocardial tissue from a COVID-19 myocarditis patient (COVID-M1). Cardiomyocytes show diffuse and disorganized cardiac troponin T (cTnT) staining with occasional cells in the blood vessel staining positively for cTnT (green). White boxes indicate zoomed in areas of the bottom figures. White arrowhead indicates cardiac troponin T material in the cytoplasm of a mononuclear cell within a blood vessel. **E**. Regions of COVID-A1 patient heart depicting normal (left) and disrupted (right) sarcomere structure. White boxes indicate zoomed in areas of the figures on the right. White arrows indicate breaks in α-actinin 2 (ACTN2) staining.

Hematoxylin & eosin staining did not reveal signs of sarcomeric damage in any samples, so we further probed the autopsy samples by immunostaining for sarcomere proteins. COVID-M1, COVID-A1, and COVID-A2 all revealed regions of severe myofibrillar anomalies. COVID-M1 displayed diffuse staining in myocytes in addition to occasional troponin-positive mononuclear cells in the vasculature **(Fig. 5D, Supplementary Fig. 6D)**, while COVID-A1 **(Fig. 5E)** and COVID-A2 **(Supplementary Fig. 6E)** displayed clear and distinct patterns of disruptions to the sarcomeric banding. We attempted to identify viral presence in the tissues using antibodies against SARS-CoV-2 Spike protein, dsRNA, and two epitopes of the viral nucleocapsid protein, but were unable to detect viral signal in any autopsy patient tissues **(Supplementary Fig. 6F)**. Overall, these patterns suggest that myocarditis in the heart leads to significant tissue damage, but is not a requirement for myofibrillar disruption of cardiac tissue. Additionally, these patterns of myocardial damage closely resemble similar cytopathic features observed by *in vitro* exposure of SARS-CoV-2 to iPSC-CMs.

## DISCUSSION

In this study, we carried out a comprehensive analysis of the cytopathic effects of SARS-CoV-2 on human iPSC-derived cardiac cell types. Cytopathic effects were particularly striking in cardiomyocytes, which manifested a distinctive myofibrillar fragmentation into individual sarcomeres and a loss of nuclear DNA from intact cell bodies. Surprisingly, these cytopathic effects appear to occur independently of the presence of actively replicating SARS-CoV-2 virus, suggesting a larger spectrum of adverse consequences than initially assumed. Guided by these observations, we observed similar sarcomeric structural disruptions and cells lacking nuclear DNA in myocardium specimens from COVID-19 patients. Together, these results provide novel insights in the pathogenesis of SARS-CoV-2 heart damage, indicate new avenues for the development of cardioprotective interventions against COVID-19, and also raise significant concerns about the prevalence and severity of cardiac involvement.

Determining the mechanisms responsible for diminished cardiac function is critically important to develop cardioprotective therapies for COVID-19. We observe that SARS-CoV-2 infection creates extremely precise and ordered disruptions to the myofibrillar structure and dissolution of the cardiac contractile machinery, which would inevitably lead to functional collapse. The striking consistency and periodicity of fragmentation suggests specific protease activity responsible for separation of the sarcomeric thick and thin filaments, as shown by immunofluorescence and TEM. Additionally, our transcriptomic analyses suggest a compensatory overexpression of myosin heavy chain genes in response to targeted degradation **(Fig 3D)**. We also observed a significant depression of the ubiquitin-proteasome system upon infection, and myofibrillar fragmentation could be partially recapitulated by the addition of the proteasome inhibitor bortezomib **(Supplementary Fig. S3; Supplementary Fig. S5G)**. Altogether, we speculate that SARS-CoV-2 infection may trigger cleavage of myofibrils by targeting myosin, which, when unopposed by the proteasome system, may result in the observed myofibrillar fragmentation phenotype.

Interestingly, myosin heavy chain protein family members contain the LKGG↓K sequence, which is known to be one of the sites used by the SARS-CoV-2 papain-like protease (PLpro) to cleave the viral polyprotein^35,36^. A cleavage of myosin by the viral protease could result in the observed filament-specific cleavage,^37^ though this interpretation does not immediately explain why myofibrillar pattern seems to be present in cells exposed to SARS-CoV-2 independently of dsRNA signal. Although further interrogation is needed to determine the precise mechanisms mediating myofibrillar fragmentation, the phenotype still informs expectations for histological specimens and therapeutic discovery.

To date, most myocardial histology from autopsy specimens of COVID-19 patients have revealed only general signs of myopathy, such as edema, occasional mononuclear infiltrate, and mild hypertrophy^18^. Although 30-50% of COVID-19 patients manifest clinical signs of cardiac dysfunction^13,38^, histological hallmarks of COVID-19 in the heart have remained elusive. Our preliminary examinations of hematoxylin and eosin staining of COVID-19 myocardial samples revealed only minor disruption and generally intact myofibrillar anatomy. However, informed by our *in vitro* analysis, we identified clear features of similar myocardial damage by immunostaining for sarcomeric proteins. This correlation reveals the potential underlying etiology of SARS-CoV-2’s impact on cardiac function and demonstrates that human iPS-derived models of myocardium can predict features of cardiac pathogenesis in COVID-19 patients.

In addition to myofibril disruption, we also identified a lack of nuclear DNA in many CMs after SARS-CoV-2 exposure, which was also observed in COVID-19 patient autopsy specimens. Physically intact cardiomyocytes lacking nuclear DNA would normally reflect a terminal event such as apoptosis^39^. However, transcriptomic analysis suggests instead that SARS-CoV-2 may be disrupting the nuclear LINC complex, through which the cytoskeleton supports the shape and structure of the nucleus^40,41^. Understanding the mechanism by which these aberrant nuclear phenotypes arise will be crucial to determine if they are a clinical feature induced by COVID-19 in the heart.

Aside from myocytes in the heart, non-myocyte cardiac stroma may also mediate some of the outcomes observed in COVID-19, such as cardiac hypertrophy and vascular dysfunction^31^. In our studies, CFs and ECs were not expected to be infectable due to their low levels of ACE2 expression. qPCR of exposed CFs and ECs supported these predictions and displayed no replicating virus, but we did observe potent cytopathic effects in both ECs and CFs. Interestingly, heat inactivated virus failed to recapitulate this effect, suggesting that abortive infection may be responsible for cytotoxicity. Future studies identifying specific mechanisms of viral toxicity to the cardiac stroma will be useful to determine how these cells may contribute to SARS-CoV-2-induced cardiac dysfunction.

The ability of iPS-CMs to model and predict cardiopathic consequences of infectious pathogens opens a wide array of potential avenues for discovery and validation of candidate cardioprotective therapies for COVID-19 and other diseases. For example, our finding that SARS-CoV-2 infects CMs via an endolysosomal route may indicate that clinical trials which target TMPRSS2 to prevent COVID-19^42,43^ may not afford effective cardioprotection without orthogonal targeting of endosomal proteases^44^. Additionally, although cell-based drug screens exist for many pathogens, including SARS-CoV-2^45^, the unique cytoarchitecture of cardiomyocytes and the specific impacts of SARS-CoV-2 on these cells offer novel screening possibilities. For example, myocyte-specific myofibrillar fragmentation enables cell-based drug screens with an exceptionally sensitive reporter of CM infection. Identification of efficacious cardioprotective therapies may require preventing viral replication and maintaining sarcomeric integrity to achieve optimal therapeutic benefit.

Although our results show that direct infection of CMs may not be required to elicit cytotoxic effects in cardiac tissue, evidence of cardiomyocytes directly infected by SARS-CoV-2 in COVID-19 patients remains scant. Analyses of the myocardium of post-mortem COVID-19 patients have detected viral dsRNA,^18^ but failed to detect staining of viral particles in cardiomyocytes at the time of death. More recent reports describe detection of SARS-CoV-2 particles for the first time inside the cardiomyocytes of COVID-19 patients^19,20^, while many autopsy specimens have undetectable virus in the heart^18,21^. We hypothesize that if virus is present in myocytes during an early to progressive stage of the disease, assays on deceased patient tissue may be limited the window of opportunity to detect transient viral presence in cardiomyocytes. However, the rapid demise in acute cases^19^ may enable observation at the peak of infection. Similarly, our *in vitro* model of CM infection allows for controlled study of the progression of the myocardial damage caused by SARS-CoV-2, thereby recapitulating earlier stages of infection.

Reports of cardiac dysfunction incidence in COVID-19 patients range from 20% to 50%, independent of disease severity^8,13,38^. Cardiac damage is strongly associated with disease mortality^3,8,18,20,46^. In addition, due to the heart’s innate lack of regenerative capacity^47,48^, a majority of these patients could suffer long-term cardiac sequelae from COVID-19. Our studies, which are analogous to mild cases of COVID-19 due to the low viral load and short time frame of viral exposure, display signs of striking cytopathic effects similar to those we observe in patient samples without any clinical diagnosis of cardiac involvement. These findings provide clues to the mechanism of cardiac pathology of COVID-19, and anticipate they will help guide the development of efficacious anti-viral and cardioprotective therapies to help manage and prevent heart damage in patients.

### Ethical Approval

Research protocol for evaluation of autopsy specimens was approved by institutional review board at the University of California, San Francisco (UCSF)/Zuckerberg San Francisco General Hospital (ZSFG).

## ACKNOWLEDGEMENTS

The following reagents were obtained through BEI Resources, NIAID, NIH: SARS-Related Coronavirus 2, Isolate USA-WA1/2020, NR-52281, deposited by the Centers for Disease Control; and Monoclonal Anti-SARS-CoV S Protein (Similar to 240C), NR-616.

We thank the Gladstone Light Microscopy and Histology Core, the Gladstone Assay Development and Drug Discovery Core, and the Gladstone Stem Cell Core for their support and experimental expertise. We also would like to thank Danielle Jorgens at the University of California Berkeley Electron Microscope for electron microscopy sample preparation and data collection. We gratefully thank the Zuckerberg-San Francisco General anatomic pathology services, including Dr. Stephen Nishimura, Mark Weinstein, and Andrew Lewis, for processing and donation of patient samples.

Endothelial cells were a kind gift from Dr. Sanjeev Ranade at the Gladstone Institutes. Patient samples were very generously contributed by Dr. Erin Brooks and Dr. Timothy Kamp at the University of Wisconsin, Madison. We thank Dr. Anita Sil, Dr. Bastian Joehnk, Dr. Lauren Rodriguez and Keith Walcott for BSL-3 laboratory support.

## Author Contributions

J.P.B., S.K., S.J.R., B.R.C. and T.C.M. designed and supervised the study. J.P.B., S.K., S.J.R., G.N.R., performed the cell culture and differentiation. C.R.S. prepared SARS-CoV-2 virus, performed the infections and all BSL3 sample collection. J.P.B., S.K., S.J.R. performed the immunostaining and imaging. J.P.B., S.K., S.J.R., G.N.R. performed the qPCR analyses. J.P.B., S.K., S.J.R., D.A.J, performed the RNA library preparation and sequencing. D.A.J. and W.R.F. performed the bioinformatic analyses. J.P.B., S.K., H.L. performed the electron microscopy studies. J.D.W. provided patient samples. A.C.S. performed the histological processing and tissue staining. All authors contributed to data analyses, writing the manuscript and figure preparation.

## METHODS

### hiPSC Maintenance; iPS-Cardiomyocyte differentiation and purification

Human iPSCs (WTc line^1^) were maintained in mTESR or mTESR+ (STEMCELL Technologies) on Matrigel (8 μg/ml, BD Biosciences)-coated cell culture plates at 37°C, 5% CO2. Cells were passaged every 3 days using Relesr (STEMCELL Technologies) and supplemented with Rock Inhibitor Y-27632 (SelleckChem) for 24 hours after each passaging. hiPSCs were differentiated into cardiomyocytes following a modified Wnt pathway modulation-based GiWi protocol^2^. Briefly, hiPSC cultures were harvested using Accutase (STEMCELL Technologies) and seeded onto Matrigel-coated 12-well plates. Three days later, cells were exposed to 12 uM CHIR99021 (Tocris) in RPMI1640 (Gibco, 11875093) supplemented with B27 without insulin (Gibco, A1895601) (R/B-) for 24 hours. After an additional 48 hours, media was changed to R/B-supplemented with 5 uM IWP2 (Tocris) for 48 hours. On day 7, media was changed to RPMI1640 medium supplemented with B27 with insulin (Gibco, 17504044) (R/B+) and refreshed every 3 days thereafter. Beating was generally observed around day 8-11. At day 15, cells were cryopreserved using CryoStor CS10 (STEMCELL Technologies). After thawing, cell cultures were enriched for iPS-cardiomyocytes following metabolic switch purification^3^. Briefly, cells were washed once with saline buffer and incubated in DMEM (without glucose, without sodium pyruvate; Gibco, 11966025) supplemented with GlutaMax (Gibco, 35050061), MEM Non-Essential Amino Acids (Gibco, 11140050) and sodium L-lactate (4mM, Sigma-Aldrich). Lactate media was refreshed every other day for a total of 6 days. Four to six days later (day 28-30), iPS-CMs were replated into assay plates for infection using 0.25% Trypsin (Gibco, 15050065) at a density of approximately 60,000 cells/cm^2^.

### scRNAseq analysis of SARS-CoV-2 entry factors

A historic single cell RNA sequencing data set consisting of iPSC-derived cardiomyocytes, primary fetal cardiac fibroblasts, and iPSC-derived endothelial cells was re-analyzed to compare relative expression levels of SARS-CoV-2 relevant receptors and proteases (GSE155226)^4^. Briefly, day 30 lactate-purified cardiomyocytes were force aggregated either alone or with a single supporting cell type and cultured in suspension culture. Aggregates were dissociated and libraries prepared using the Chromium 3’ v2 library preparation platform (10X Genomics). Libraries were sequenced on a NextSeq 550 sequencer (Illumina) to a depth of at least 30 million reads per sample. Samples were demultiplexed and aligned to GRCh38 with CellRanger v3.0.2. Individual cell UMIs were filtered using Seurat v3.2.0^5^, keeping only cells with at least 1,000 reads, 300 detected genes, and less than 10% mitochondrial reads. The top 2,000 variable genes were projected onto 20 principal components. Although greater than 5% of cells were detected in either S or G2M phase, regressing out cell cycle genes did not alter clustering of primary cell types. Cells were clustered with a resolution of 0.4, yielding three primary clusters corresponding to each cell type, which were used to profile cell-type specific expression of SARS-CoV-2 relevant factors.

### Cardiac Fibroblast Differentiation

Second heart field-derived cardiac fibroblasts (SHF-CFs) were differentiated following the GiFGF protocol, as previously published^6^. Briefly, hiPSCs were seeded at 15,000 cells/cm^2^ in mTeSR1 medium. Once they reached 100% confluency, they were treated with R/B-supplemented with 12μM CHIR99021 (day 0), and refreshed with R/B-24 hours later (day 1). From days 2-20, cells were fed every 2 days with cardiac fibroblast basal media (CFBM) (Lonza, CC-3131) supplemented with 75ng/mL bFGF. On day 20, CFs were singularized with Accutase for 10 minutes and replated at 7,000 cells/cm^2^ onto tissue culture plastic 10cm dishes in FibroGRO medium (Millipore Sigma, SCMF001). FibroGRO media was changed every two days until the CFs reached approximately 80-90% confluency, at which point they were passaged with Accutase. SHF-CFs were validated to be >80% double-positive for TE-7 and vimentin by flow cytometry.

### Endothelial Cell Differentiation

WTC11 iPSCs were directed towards an endothelial cell (EC) lineage by the addition of E8 media supplemented with BMP4 (5 ng/ml) and Activin A (25 ng/ml) for 48 hours followed by E7BVi media, consisting of E6 medium supplemented with bFGF (50ng/ml), VEGF-A (50 ng/ml), BMP4 (50 ng/ml) and a TGFβ inhibitor, SB431542, (5 μM) for 72 hours. After 5 days of successive media changes, ECs were split and plated at high density in EGM media (Lonza, CC-3162) on tissue culture flasks coated with fibronectin (1:100, Sigma Aldrich F0895). On day 8, all cells were cryo-preserved and a fraction of ECs were assayed for >95% purity by flow cytometry using antibodies against mature EC markers CD31 and CDH5. ECs were seeded at a density of roughly 60,000 cells/cm^2^, 48 hours prior to infection.

### Mixed Cultures of CMs, CFs, and ECs

Mixed cultures of iPS-CMs, iPS-ECs, and iPS-CFs were created by combining single cell suspensions of each cell types in a ratio of 60:30:10 CM:EC:CF at a density of 200,000 cells/mL. The mixed suspension was replated onto Matrigel-coated tissue culture plates 48 hours prior to infection at a density of 62,500 cells/cm^2^.

### SARS-CoV-2 Infection

The WA-1 strain (BEI resources) of SARS-CoV-2 was used for all experiments. All live virus experiments were performed in a Biosafety Level 3 lab. SARS-CoV-2 stocks were passaged in Vero cells (ATCC) and titer was determined via plaque assay on Vero cells as previously described^7^. Briefly, virus was diluted 1:10^2^-1:10^6^ and incubated for 1 hour on Vero cells before an overlay of Avicel and complete DMEM (Sigma Aldrich, SLM-241) was added. After incubation at 37°C for 72 hours, the overlay was removed and cells were fixed with 10% formalin, stained with crystal violet, and counted for plaque formation. SARS-CoV-2 infections of iPSc and iPS-derived cardiac cells were done at a multiplicity of infection of 0.006 for 48 hours unless otherwise specified. For heat inactivation, SARS-CoV-2 stocks were incubated at 85°C for 5 min. Plaque assay for supernatant from infected cells was performed as above.

### Immunocytochemistry

Infected and mock-treated cell cultures in coverslips were washed with Phosphate Buffered Solution (PBS) and fixed in 4% paraformaldehyde (PFA) overnight, followed by blocking and permeabilization with 0.1% Triton-X 100 (T8787, Sigma) and 5% BSA (A4503, Sigma) for one hour at RT. Antibody dilution buffer (Ab buffer) was comprised of PBS supplemented with 0.1% Triton-X 100 and 1% BSA. Samples were incubated with primary antibodies overnight at 4°C **(Table 2)**, followed by 3 washes with PBS and incubation with fluorescent-conjugated secondary antibodies at 1:250 in Ab buffer for 1 hour at RT **(Table 2)**. Coverslips were mounted onto SuperFrost Slides (FisherBrand, 12-550-15) with ProLong Antifade mounting solution with DAPI (Invitrogen, P36931. Images were acquired with a Zeiss Axio Observer Z.1 Spinning Disk Confocal (Carl Zeiss) or with an ImageXpress Micro Confocal High-Content Imaging System (Molecular Devices) and processed using ZenBlue and ImageJ.

**Table 1:** List of reagents.

| Drug | Concentration | Provider |
| --- | --- | --- |
| DMSO | 0.1% (1:1000) | Fisher Scientific (BP231-100) |
| Interferon $\alpha$ | 2500U/mL | Sigma Aldrich (SRP4596-100UG) |
| Interferon $\beta$ | 2500U/mL | Sigma Aldrich (IF014) |
| Interferon $\gamma$ | 2500U/mL | BioRad (PHP050A) |
| Interferon $\lambda$ | 2500U/mL | Cedarlane Laboratories (CLY100-169-5UG) |
| Ruxolitinib | 500nM | Thermo (NC1399519) |
| Doxorubicin | 20nM and 200nM | Sigma (D1515) |
| Bortezomib | 1uM and 10uM | Sigma (CAS 179324-69-7) |
| Bafilomycin | 100nM | Cayman Chemical (98813-13-9) |
| Aprotinin | 50uM | Cayman Chemical (9087-70-1) |
| Camostat mesilate | 2uM | Cayman Chemical (59721-29-8) |
| CA074 | 30uM | Cayman Chemical (134448-10-5) |
| E-64d | 25uM | Cayman Chemical (88321-09-9) |
| Z-Phe-Tyr(tBu)-<br>diazomethylketone | 30uM | Cayman Chemical (114014-15-2) |
| ACE2 blocking antibody | 20ug/ml | R&D Systems (AF933) |
| Apilimod | 1uM | SelleckChem Chemicals (S6414) |

**Table 2:**
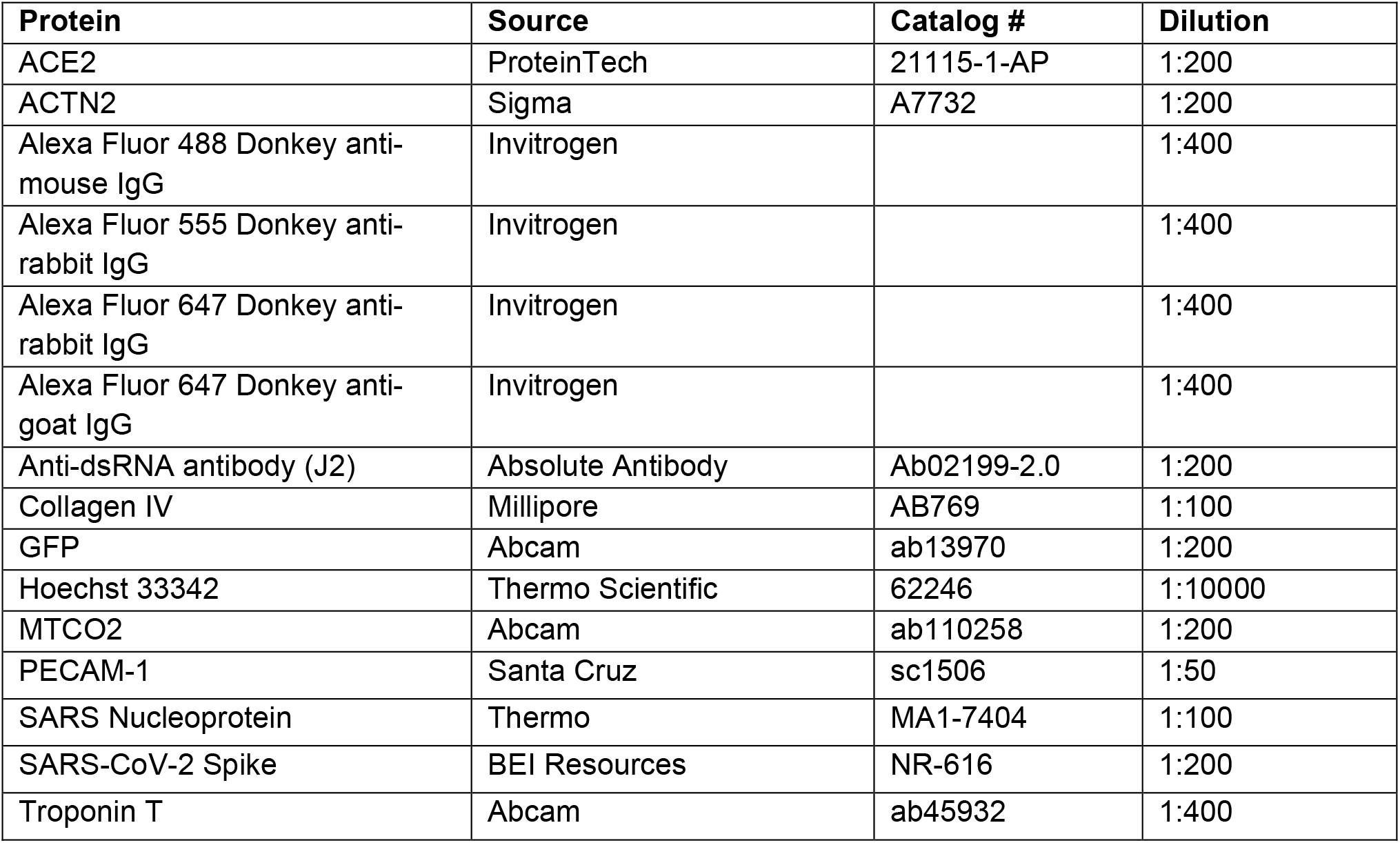
List of dyes, primary and secondary antibodies used for immunocytochemistry and paraffin sections.

| Protein | Source | Catalog # | Dilution |
| --- | --- | --- | --- |
| ACE2 | ProteinTech | 21115-1-AP | 1:200 |
| ACTN2 | Sigma | A7732 | 1:200 |
| Alexa Fluor 488 Donkey anti-mouse IgG | Invitrogen |  | 1:400 |
| Alexa Fluor 555 Donkey anti-rabbit IgG | Invitrogen |  | 1:400 |
| Alexa Fluor 647 Donkey anti-rabbit IgG | Invitrogen |  | 1:400 |
| Alexa Fluor 647 Donkey anti-goat IgG | Invitrogen |  | 1:400 |
| Anti-dsRNA antibody (J2) | Absolute Antibody | Ab02199-2.0 | 1:200 |
| Collagen IV | Millipore | AB769 | 1:100 |
| GFP | Abcam | ab13970 | 1:200 |
| Hoechst 33342 | Thermo Scientific | 62246 | 1:10000 |
| MTCO2 | Abcam | ab110258 | 1:200 |
| PECAM-1 | Santa Cruz | sc1506 | 1:50 |
| SARS Nucleoprotein | Thermo | MA1-7404 | 1:100 |
| SARS-CoV-2 Spike | BEI Resources | NR-616 | 1:200 |
| Troponin T | Abcam | ab45932 | 1:400 |

**Table 3:**
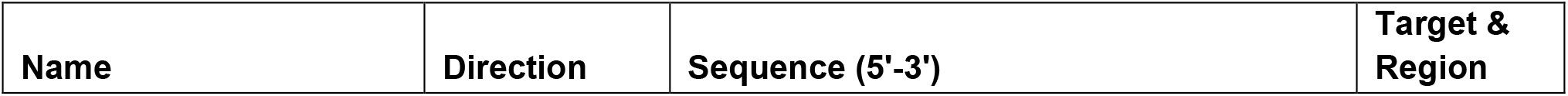
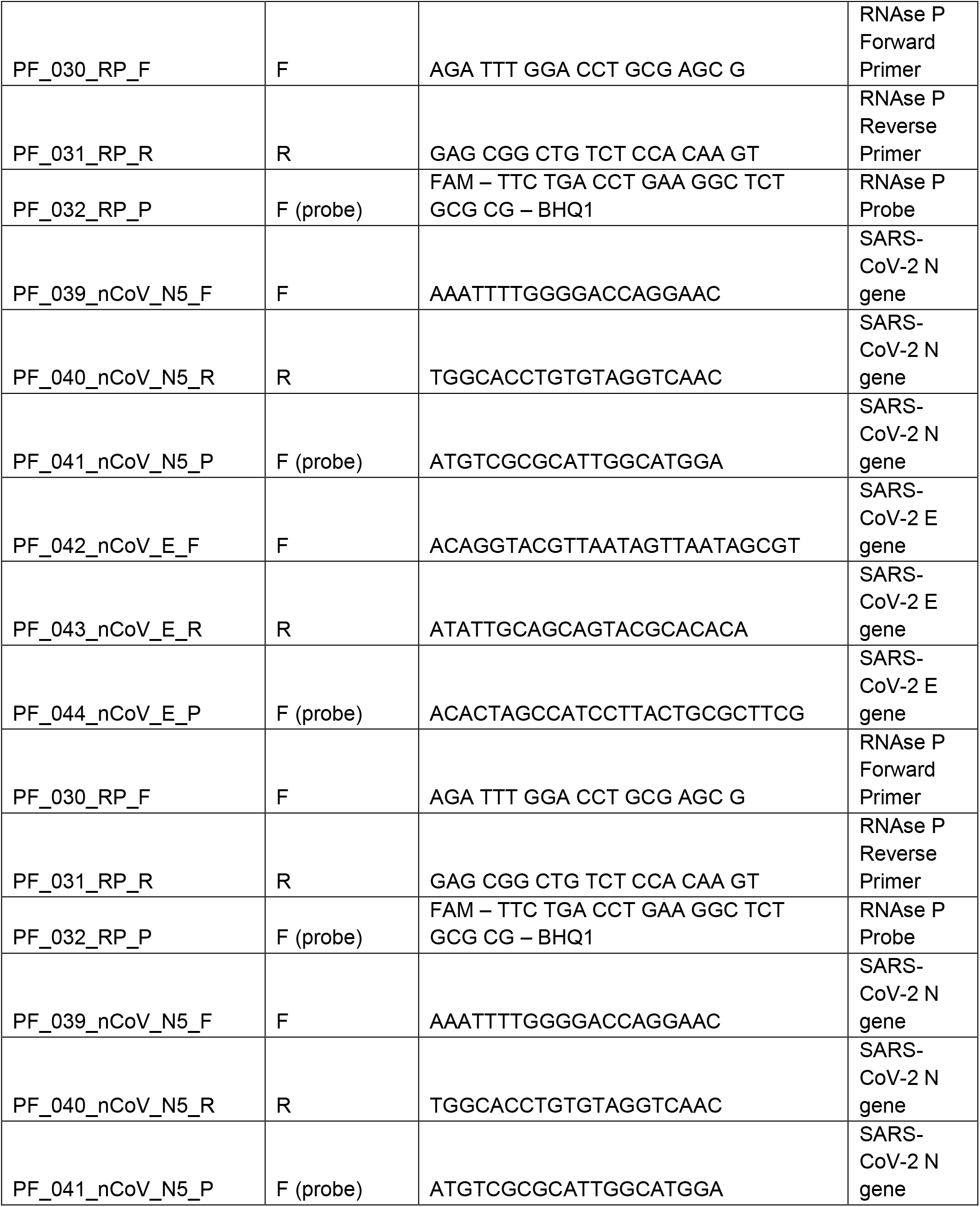
List of primers.

### Histology

Paraffin section of healthy and COVID-19 patient hearts were deparaffinized using xylene, re-hydrated through a series of decreasing ethanol solutions (100%, 100%, 95%, 80%, 70%) and rinsed in PBS. Hematoxylin and eosin staining was performed according manufacturer instructions and the slides were mounted with Cytoseal 60 (Richard-Allan Scientific) and glass coverslips. For immunofluorescence staining, epitopes were retrieved by immersing slides through 35 min incubation at 95°C in citrate buffer (Vector Laboratories, pH 6) or Tris-EDTA buffer (Cellgro, pH 9). Slides were cooled for 20min at RT and washed with PBS. Samples were permeabilized in 0.2% Triton X-100 (Sigma) in PBS by slide immersion and washed in PBS.

Blocking was performed in 1.5% normal donkey serum (NDS; Jackson ImmunoResearch) and PBS solution for 1h at RT. Primary and secondary antibody cocktails were diluted in blocking solution **(Table 2)**. PBS washes were performed after primary (overnight, 4°C) and secondary antibody (1h, RT) incubations. Nuclei were stained with Hoechst and coverslips were mounted on slides using ProLong™ Gold Antifade Mountant. Samples were imaged on the Zeiss Axio Observer Z1.

### RT-qPCR

Cultured cells were lysed with Qiagen buffer RLT (Qiagen, 79216) supplemented with 1% β-mercaptoethanol (Bio-Rad, 1610710) and RNA was isolated using the RNeasy Mini Kit (Qiagen 74104) or Quick-RNA MicroPrep (ThermoFisher, 50444593) and quantified using the NanoDrop 2000c (ThermoFisher). Viral load was measured by detection of the viral Nucleocapsid (N) transcript through one-step quantitative real-time PCR, performed using PrimeTime Gene Expression Master Mix (Integrated DNA Technologies, 1055772) with primers and probes specific to N (‘N5’) and RPP30 as in internal reference. RT-qPCR reactions were performed on a CFX384 (BioRad) and delta cycle threshold (ΔCt) was determined relative to RPP30 levels. Viral detection levels in pharmacologically treated samples were normalized to DMSO-treated controls.

### RNA-Seq

To generate libraries for RNA-sequencing, RNA isolate quality was assessed with an Agilent Bioanalyzer 2100 on using the RNA Pico Kit (Agilent, 5067-1513). 10ng of each RNA isolate was then prepared using the Takara SMARTer Stranded Total RNA-Seq Kit v2 – Pico Input Mammalian (Takara, 634412). Transcripts were fragmented for 3.5 minutes and amplified for 12 cycles. Library concentrations were quantified with the Qubit dsDNA HS Assay Kit (Thermo Fisher, Q32851) and pooled for sequencing. Sequencing was performed on an Illumina NextSeq 550 system, using the NextSeq 500/550 High Output Kit v2.5 (150 Cycles) (Illumina, 20024907) to a depth of at least 10 million reads per sample. Raw data is available at GEO under the accession number GSE156754.

### Bioinformatic analyses of transcriptomic data

Samples were demultiplexed using bcl2fastq v2.20.0 and aligned to both GRCh38 and the SARS-CoV-2 reference sequence (NC_045512) using hisat2 v2.1.0^8^. Aligned reads were converted to counts using featureCounts v1.6.2^9^. Cell-type clustering, gene loadings, and technical replication were assessed using the PCA and MDS projections implemented in scikit-learn v0.23^10^. Aside from one outlier fibroblast sample with low read depth, two different batches of fibroblasts and endothelial cells clustered together in both PCA and MDS space, suggesting minimal batch to batch variation. Only one batch of CMs and iPSCs were sequenced per condition, so batch to batch variability could not be assessed for those cell types. Differential expression analysis was performed using edgeR v3.30.2^11^ with limma/voom v3.44.3 normalization^12^ and GO term enrichment analysis was performed using clusterProfiler v3.16.0^13^. Correlation of gene expression with distance in PCA space from mock CMs was assessed using Pearson’s R test. Pathways for sarcomere organization and the LINC complex were adapted from WikiPathways (WP383 and WP4535 respectively) using Cytoscape v2.8.0^14^.

### TEM/CLEM

Cells grown on gridded 35mm MatTek glass-bottom dishes (MatTek Corp., Ashland, MA, USA) were fixed in 2.5% glutaraldehyde and 2.5% paraformaldehyde in 0.1M sodium cacodylate buffer, pH 7.4 (EMS, Hatfield, PA, USA) following fluorescence imaging. Samples were rinsed 3 x 5 min at RT in 0.1M sodium cacodylate buffer, pH 7.2, and immersed in 1% osmium tetroxide with 1.6% potassium ferricyanide in 0.1M sodium cacodylate buffer for 30 minutes. Samples were rinsed (3 x 5 min, RT) in buffer and briefly washed with distilled water (1 x 1 min, RT) before sample were then subjected to an ascending ethanol gradient (7 min; 35%, 50%, 70%, 80%, 90%) followed by pure ethanol. Samples were progressively infiltrated (using ethanol as the solvent) with Epon resin (EMS, Hatfield, PA, USA) and polymerized at 60°C for 24-48 hours. Care was taken to ensure only a thin amount of resin remained within the glass bottom dishes to enable the best possible chance for separation of the glass coverslip. Following polymerization, the glass coverslips were removed using ultra-thin Personna razor blades (EMS, Hatfield, PA, USA) and liquid nitrogen exposure as needed. The regions of interest, identified by the gridded alpha-numerical labeling, were carefully removed and mounted with cyanoacrylate glue for sectioning on a blank block. Serial thin sections (100 nm) were cut using a Leica UC 6 ultramicrotome (Leica, Wetzlar, Germany) from the surface of the block until approximately 4-5 microns in to ensure complete capture of the cell volumes. Section-ribbons were then collected sequentially onto formvar-coated 50 mesh copper grids. The grids were post-stained with 2% uranyl acetate followed by Reynold’s lead citrate, for 5 min each. The sections were imaged using a Tecnai 12 120kV TEM (FEI, Hillsboro, OR, USA), data recorded using an UltraScan 1000 with Digital Micrograph 3 software (Gatan Inc., Pleasanton, CA, USA), and montaged datasets were collected with SerialEM (bio3d.colorado.edu/SerialEM) and reconstructed using IMOD eTOMO (bio3d.colorado.edu/imod). Reconstructed images were denoised using total variational denoising, followed by gamma correction and unsharp masking (https://peerj.com/articles/453/).

### Analysis of immunofluorescence images

Immunostained images of cells were coded and manually counted by four blinded individuals, with a 20% overlap for concordance. Features counted were dsRNA positive cells (defined as a cell with clear dsRNA staining in the cytoplasm over background), cells presenting myofibrillar fragmentation (defined as presenting at least one instance of a cTnT positive doublet obviously separated from other myofibrils), and cells positive for both dsRNA and myofibrillar fragmentation. Each datapoint represents the normalized sum of counts for nine randomly acquired fields of view in a separate well using high magnification (40x). Counts were normalized to total nuclei counts per field of view. For viability estimations, each data point is the sum of nine randomly acquired fields of view in a separate well, using low magnification (10x). Nuclei counts were performed automatically using the EBImage package^15^ on R^16^. For the analysis COVID-19 patient samples, nuclei counts were performed manually, and statistical analysis was done by fit to a Poisson generalized linear model.

## Statistical Analyses

Statistical testing for qPCR experiments was performed using GraphPad Prism 8 software, using 1-way ANOVA with post-hoc Tukey’s multiple comparisons test. Statistical analysis for the immunofluorescence cell counts was performed using R^16^ using Student’s t-test with Bonferroni correction for multiple testing. Statistical differences in expression between bioinformatic samples were performed on corrected, log transformed counts using Welch’s t-test with Benjamini Hochberg false discovery rate correction with a threshold of 0.05, except where described above.

