## Supplementary Information for "SARS-CoV-2 infection of human iPSC-derived cardiac cells predicts novel cytopathic features in hearts of COVID-19 patients"

### SUPPLEMENTARY FIGURES

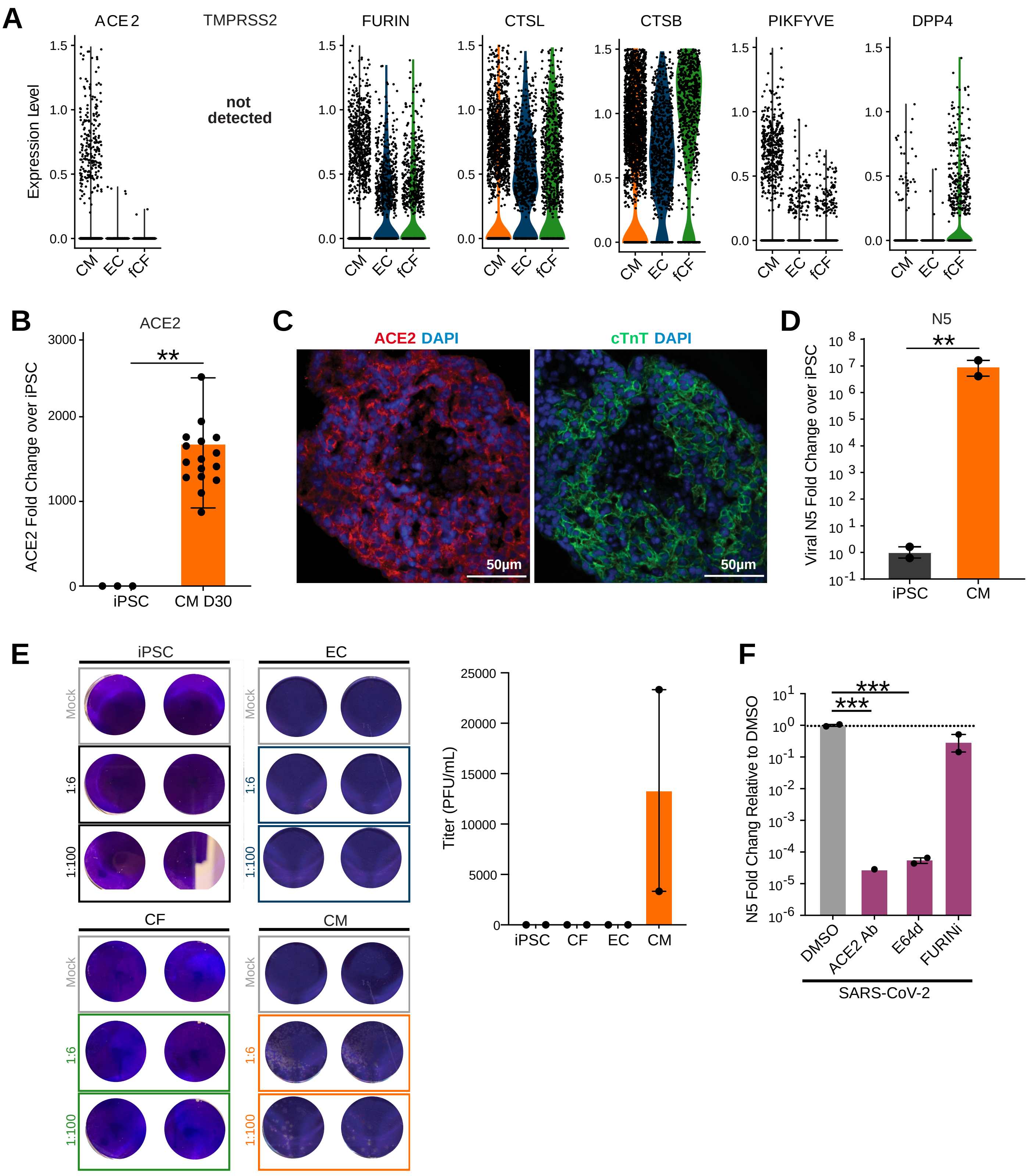

**Supplementary Figure 1. SARS-CoV-2 in iPSC-derived cardiac cells, RNA levels of entry factors, productive infection assay and drug pretreatment effects. A.** Single-cell transcript levels for factors of interest. fCF: fetal cardiac fibroblast. Each dot represents normalized transcript levels in a single cell. **B.** RT-qPCR quantification of ACE2 transcript levels in 30-day CMs compared to undifferentiated iPSCs. **: p-value < 0.01. **C.** Representative immunostaining of mixed cardiac cell aggregates. Two sequential sections are shown, highlighting ACE2 receptor and cardiac troponin T (cTnT) staining. **D.** Infection of iPSCs and CMs yields no detectable infection in iPSCs. **E.** Images of plaque assay from supernatant from SARS-CoV-2-infected cardiac cell types. Crystal Violet stains are shown for serial dilutions of biological duplicate samples. Bar graph provides quantification of plaque forming units in supernatant. **F.** RT-qPCR quantification of SARS-CoV-2 RNA levels in CMs pretreated with different viral infection blocking agents. Graph depicts fold change relative to vehicle control (DMSO). Duplicates were analyzed for significance by one-way ANOVA with Tukey's multiple comparisons. ***: p-value < 0.001.

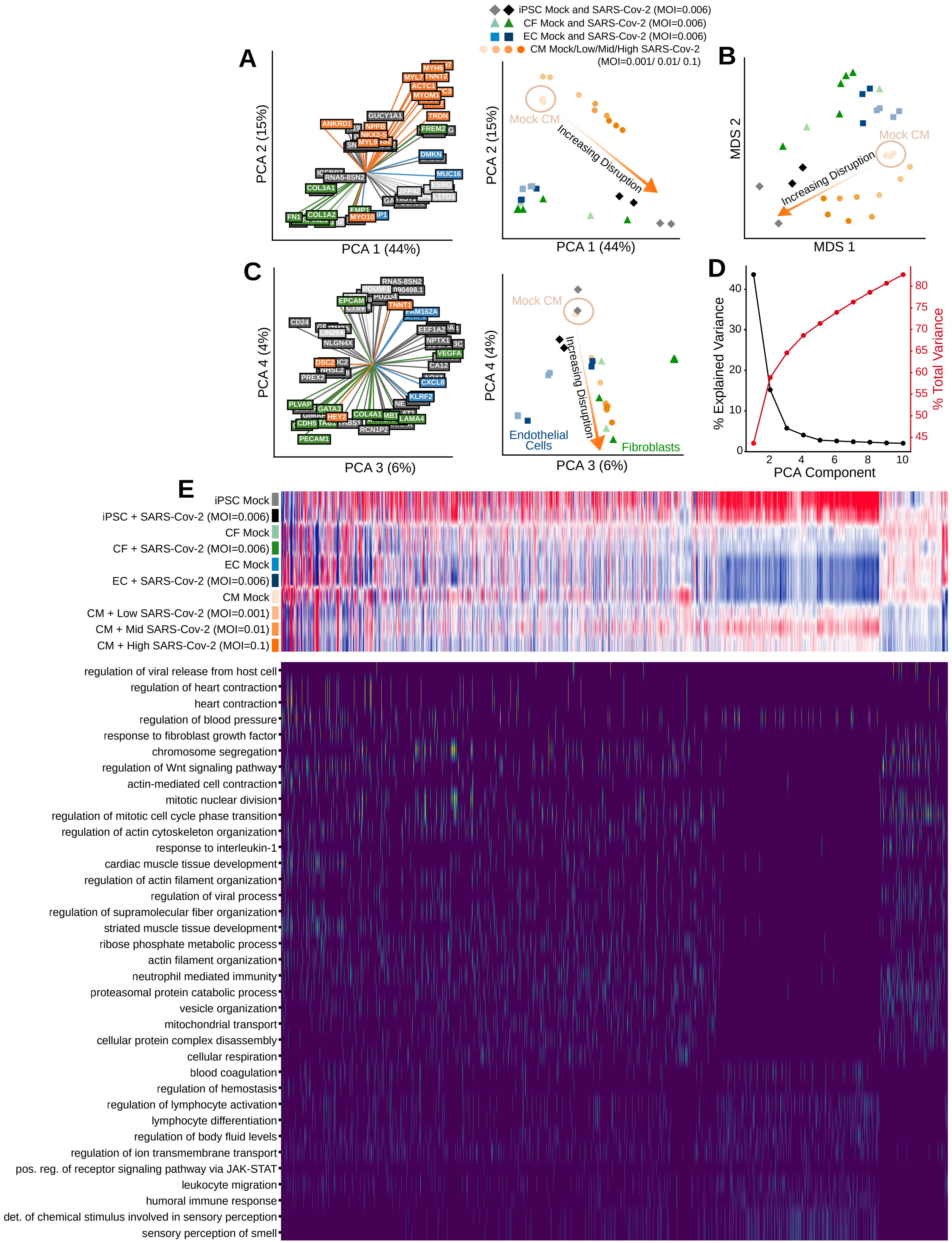
iPSC Mock and SARS-Cov-2 (MOI=0.006)
CF Mock and SARS-Cov-2 (MOI=0.006)
EC Mock and SARS-Cov-2 (MOI=0.006)
CM Mock/Low/Mid/High SARS-Cov-2
(MOI=0.001/ 0.01/ 0.1)
A
B
C
D
E
PCA 1 (44%)
PCA 2 (15%)
Mock CM
Increasing Disruption
MDS 1
MDS 2
PCA 3 (6%)
PCA 4 (4%)
Endothelial
Cells
Fibroblasts
% Explained Variance
% Total Variance
PCA Component
iPSC Mock
iPSC + SARS-Cov-2 (MOI=0.006)
CF Mock
CF + SARS-Cov-2 (MOI=0.006)
EC Mock
EC + SARS-Cov-2 (MOI=0.006)
CM Mock
CM + Low SARS-Cov-2 (MOI=0.001)
CM + Mid SARS-Cov-2 (MOI=0.01)
CM + High SARS-Cov-2 (MOI=0.1)
regulation of viral release from host cell
regulation of heart contraction
heart contraction
regulation of blood pressure
response to fibroblast growth factor
chromosome segregation
regulation of Wnt signaling pathway
actin-mediated cell contraction
mitotic nuclear division
regulation of mitotic cell cycle phase transition
regulation of actin cytoskeleton organization
response to interleukin-1
cardiac muscle tissue development
regulation of actin filament organization
regulation of viral process
regulation of supramolecular fiber organization
striated muscle tissue development
ribose phosphate metabolic process
actin filament organization
neutrophil mediated immunity
proteasomal protein catabolic process
vesicle organization
mitochondrial transport
cellular protein complex disassembly
cellular respiration
blood coagulation
regulation of hemostasis
regulation of lymphocyte activation
lymphocyte differentiation
regulation of body fluid levels
regulation of ion transmembrane transport
pos. reg. of receptor signaling pathway via JAK-STAT
leukocyte migration
humoral immune response
det. of chemical stimulus involved in sensory perception
sensory perception of smell

**Supplementary Figure 2. Transcriptional disruption and cellular pathway dysregulation due to SARS-CoV-2 exposure in cardiac cells. A.** Loading plot of top genes expressed along radial combinations of principal components 1 and 2 with location of iPSCs, CFs, ECs, and CMs ranked by PCA distance from the mock CM cluster and gene color indicating cardiomyocyte state (orange), fibroblast/endothelial cell state (green), iPSC state (light gray), immune response (blue), or other function (dark gray). **B.** Multi-dimensional scaling (MDS) projection of iPSCs, CFs, ECs, and CMs ranked by MDS distance from the mock CM cluster. **C.** Loading plot of top genes expressed along radial combinations of principal components 3 and 4 with location of iPSCs, CFs, ECs, and CMs ranked by PCA distance from the mock CM cluster and gene color indicating cardiomyocyte state (orange), fibroblast/endothelial cell state (green), iPSC state (light gray), immune response (blue), or other function (dark gray). **D.** Percent variance explained (black) and cumulative variance explained (red) by the first 10 principal components. **E.** Heat map depicting transcriptional expression differences between iPSCs and different cardiac cell types in both mock and SARS-CoV-2 infected treatments, with CMs exposed to different MOIs. Genes map to GO terms of interest (Genes |log2 fold change| > 1 between high infection and mock, FDR < 0.05; GO terms containing at least 25 enriched genes, FDR < 0.01).

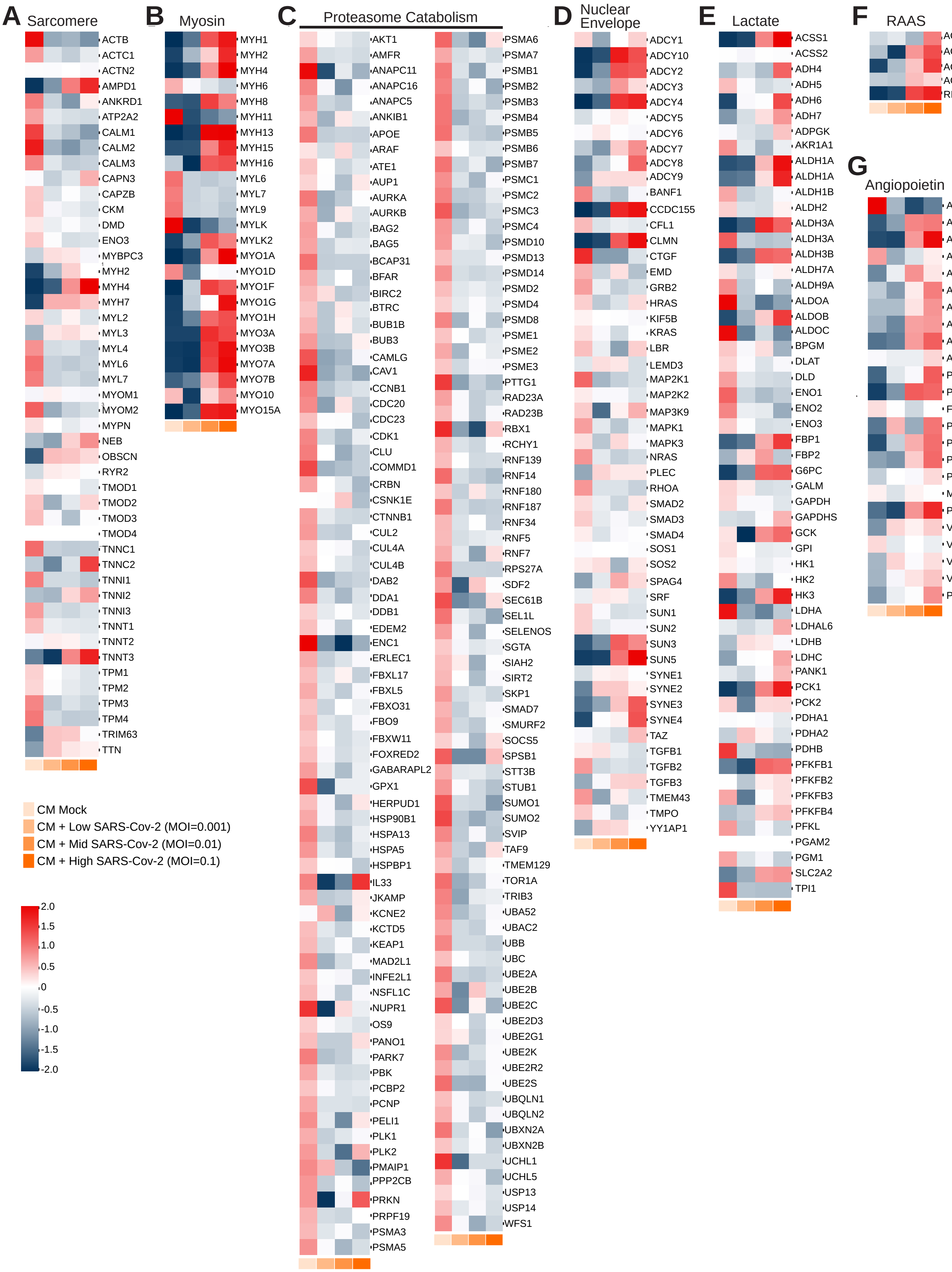
A Sarcomere
ACTB
ACTC1
ACTN2
AMPD1
ANKRD1
ATP2A2
CALM1
CALM2
CALM3
CAPN3
CAPZB
CKM
DMD
ENO3
MYBPC3
MYH2
MYH4
MYH7
MYL2
MYL3
MYL4
MYL6
MYL7
MYOM1
MYOM2
MYPN
NEB
OBSCN
RYR2
TMOD1
TMOD2
TMOD3
TMOD4
TNNC1
TNNC2
TNNI1
TNNI2
TNNI3
TNNT1
TNNT2
TNNT3
TPM1
TPM2
TPM3
TPM4
TRIM63
TTN
B Myosin
MYH1
MYH2
MYH4
MYH6
MYH8
MYH11
MYH13
MYH15
MYH16
MYL6
MYL7
MYL9
MYLK
MYLK2
MYO1A
MYO1D
MYO1F
MYO1G
MYO1H
MYO3A
MYO3B
MYO7A
MYO7B
MYO10
MYO15A
C Proteasome Catabolism
AKT1
AMFR
ANAPC11
ANAPC16
ANAPC5
ANKIB1
APOE
ARAF
ATE1
AUP1
AURKA
AURKB
BAG2
BAG5
BCAP31
BFAR
BIRC2
BTRC
BUB1B
BUB3
CAMLG
CAV1
CCNB1
CDC20
CDC23
CDK1
CLU
COMMD1
CRBN
CSNK1E
CTNNB1
CUL2
CUL4A
CUL4B
DAB2
DDA1
DDB1
EDEM2
ENC1
ERLEC1
FBXL17
FBXL5
FBXO31
FBO9
FBXW11
FOXRED2
GABARAPL2
GPX1
HERPUD1
HSP90B1
HSPA13
HSPA5
HSPBP1
IL33
JKAMP
KCNE2
KCTD5
KEAP1
MAD2L1
INFE2L1
NSFL1C
NUPR1
OS9
PANO1
PARK7
PBK
PCBP2
PCNP
PELI1
PLK1
PLK2
PMAIP1
PPP2CB
PRKN
PRPF19
PSMA3
PSMA5
PSMA6
PSMA7
PSMB1
PSMB2
PSMB3
PSMB4
PSMB5
PSMB6
PSMB7
PSMC1
PSMC2
PSMC3
PSMC4
PSMD10
PSMD13
PSMD14
PSMD2
PSMD4
PSMD8
PSME1
PSME2
PSME3
PTTG1
RAD23A
RAD23B
RBX1
RCHY1
RNF139
RNF14
RNF180
RNF187
RNF34
RNF5
RNF7
RPS27A
SDF2
SEC61B
SEL1L
SELENOS
SGTA
SIAH2
SIRT2
SKP1
SMAD7
SMURF2
SOCS5
SPSB1
STT3B
STUB1
SUMO1
SUMO2
SVIP
TAF9
TMEM129
TOR1A
TRIB3
UBA52
UBAC2
UBB
UBC
UBE2A
UBE2B
UBE2C
UBE2D3
UBE2G1
UBE2K
UBE2R2
UBE2S
UBQLN1
UBQLN2
UBXN2A
UBXN2B
UCHL1
UCHL5
USP13
USP14
WFS1
D Nuclear Envelope
ADCY1
ADCY10
ADCY2
ADCY3
ADCY4
ADCY5
ADCY6
ADCY7
ADCY8
ADCY9
BANF1
CCDC155
CFL1
CLMN
CTGF
EMD
GRB2
HRAS
KIF5B
KRAS
LBR
LEMD3
MAP2K1
MAP2K2
MAP3K9
MAPK1
MAPK3
NRAS
PLEC
RHOA
SMAD2
SMAD3
SMAD4
SOS1
SOS2
SPAG4
SRF
SUN1
SUN2
SUN3
SUN5
SYNE1
SYNE2
SYNE3
SYNE4
TAZ
TGFB1
TGFB2
TGFB3
TMEM43
TMPO
YY1AP1
E Lactate
ACSS1
ACSS2
ADH4
ADH5
ADH6
ADH7
ADPGK
AKR1A1
ALDH1A
ALDH1A
ALDH1B
ALDH2
ALDH3A
ALDH3A
ALDH3B
ALDH7A
ALDH9A
ALDOA
ALDOB
ALDOC
BPGM
DLAT
DLD
ENO1
ENO2
ENO3
FBP1
FBP2
G6PC
GALM
GAPDH
GAPDHS
GCK
GPI
HK1
HK2
HK3
LDHA
LDHAL6
LDHB
LDHC
PANK1
PCK1
PCK2
PDHA1
PDHA2
PDHB
PFKFB1
PFKFB2
PFKFB3
PFKFB4
PFKL
PGAM2
PGM1
SLC2A2
TPI1
F RAAS
ACE2
ACE
AGTR1
AGTR2
REN
G Angiopoietin
ANGPT1
ANGPT2
ANGPT4
ANGPTL2
ANGPTL3
ANGPTL4
ANGPTL5
ANGPTL6
ANGPTL7
ANGPTL8
PCSK1
PCSK2
FURIN
PCSK4
PCSK5
PCSK6
PCSK7
MBTPS1
PCSK9
VEGFA
VEGFB
VEGFC
VEGFD
PGF
CM Mock
CM + Low SARS-Cov-2 (MOI=0.001)
CM + Mid SARS-Cov-2 (MOI=0.01)
CM + High SARS-Cov-2 (MOI=0.1)
2.0
1.5
1.0
0.5
0
-0.5
-1.0
-1.5
-2.0

**Supplementary Figure 3.** Heat maps for gene pathways of interest, comparing transcriptional differences in mock and SARS-CoV-2-infected cardiomyocytes. Pathway schematics for each pathway showing individual genes upregulated or downregulated in infection (red and blue respectively). Gene sets display notable changes in regulation of (**A**) sarcomere structure, (**B**) myosin contractility, (**C**) proteasome catabolic processes, (**D)** nuclear envelope, (**E**) lactate metabolism, (**F)** renin-angiotensin pathways, and (**G**) angiopoietin pathway.

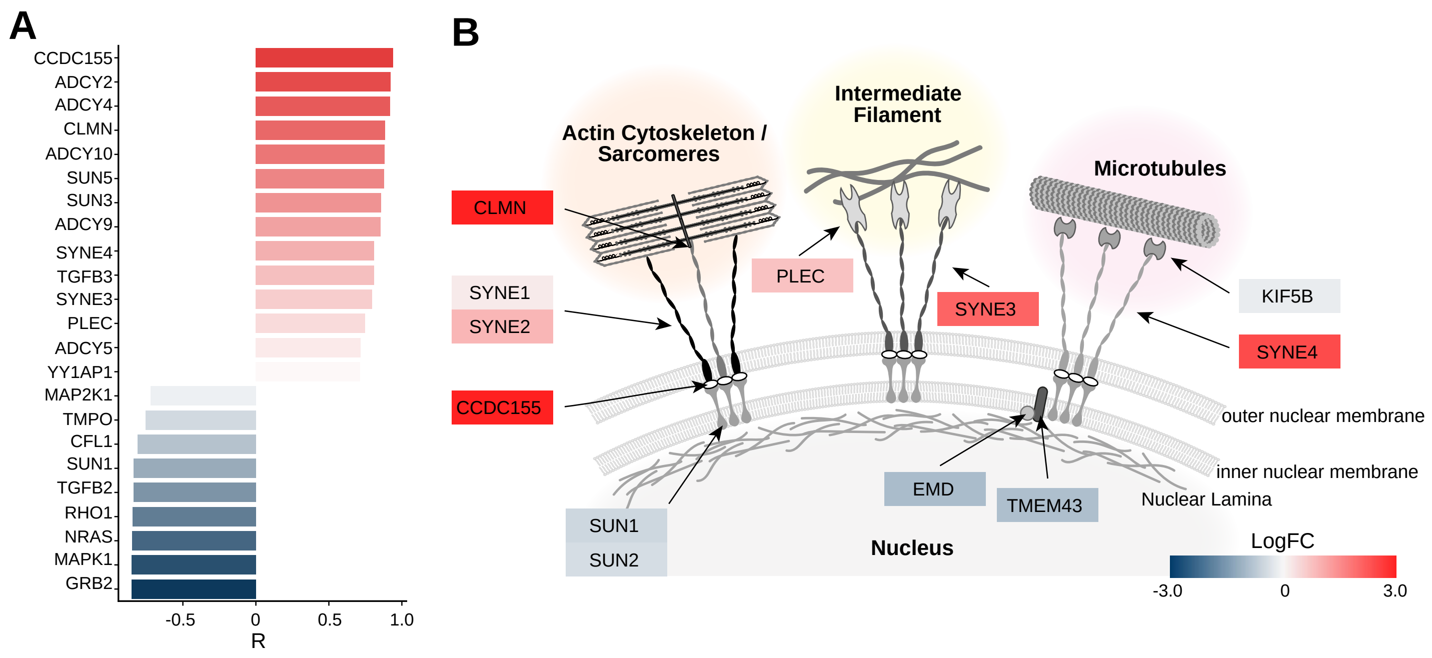


**Supplementary Figure 4. A.** Expression ratio of genes involved in nuclear envelope structure and stabilization between the high infection and mock infection CM groups. **B.** Schematic of the nuclear envelope showing localization of differentially regulated factors, with dysregulation of SYNE3/4 and SUN1/2. Background colors denote log2FC of the high infected condition over mock.

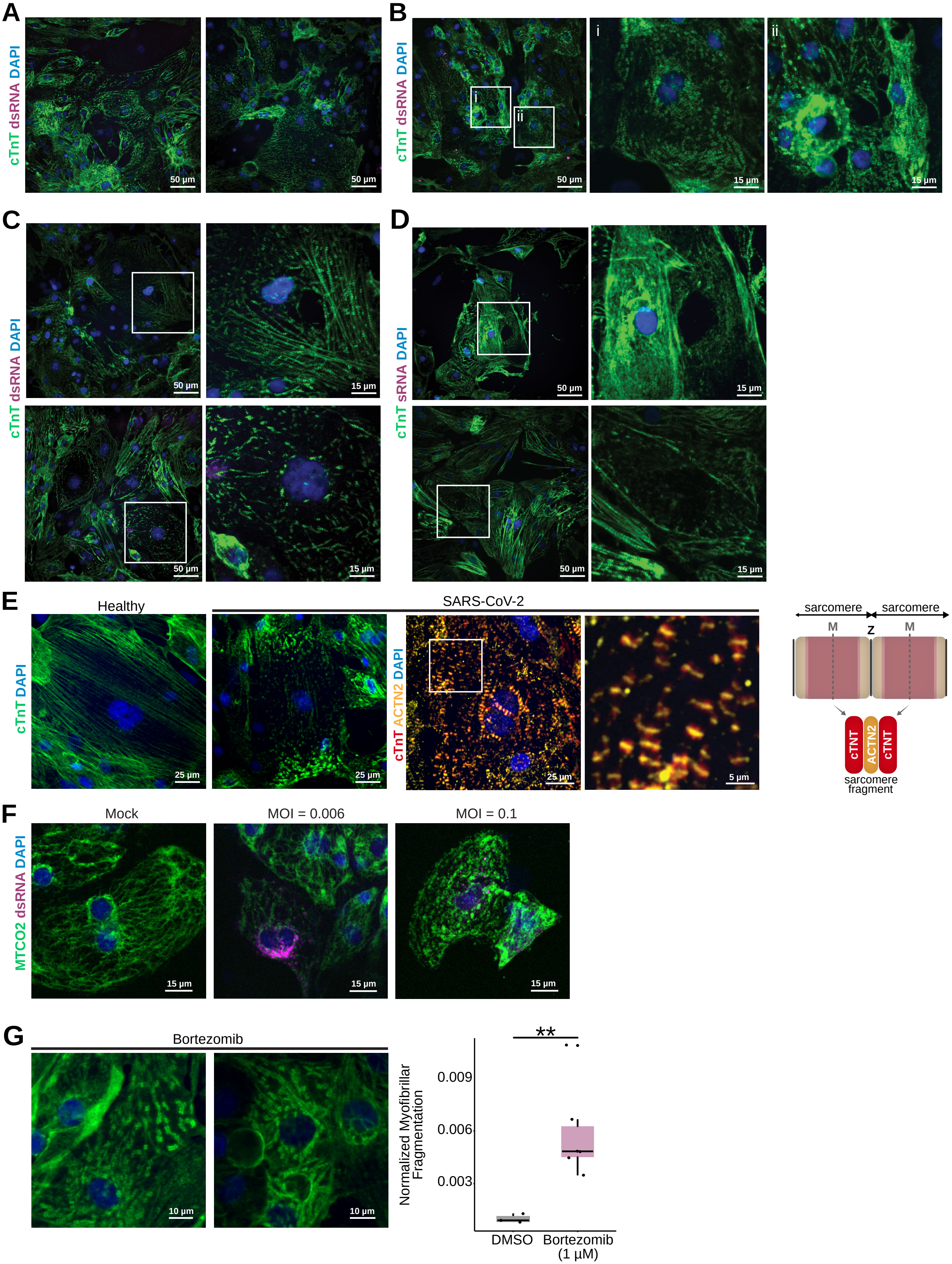
A
cTnT dsRNA DAPI
50 µm
50 µm
B
cTnT dsRNA DAPI
i
ii
50 µm
15 µm
15 µm
C
cTnT dsRNA DAPI
50 µm
15 µm
50 µm
15 µm
D
cTnT sRNA DAPI
50 µm
15 µm
50 µm
15 µm
E
Healthy
SARS-CoV-2
cTnT DAPI
cTnT ACTN2 DAPI
25 µm
25 µm
25 µm
5 µm
sarcomere
sarcomere
M
Z
M
cTNT
ACTN2
cTNT
sarcomere fragment
F
Mock
MOI = 0.006
MOI = 0.1
MTCO2 dsRNA DAPI
15 µm
15 µm
15 µm
G
Bortezomib
10 µm
10 µm
**
Normalized Myofibrillar Fragmentation
0.009
0.006
0.003
DMSO
Bortezomib (1 µM)

**Supplementary Figure 5. A-D.** Additional representative images of myofibrillar fragmentation observed in SARS-CoV-2-exposed CMs, highlighting (**A**) the prevalence of myofibrillar fragmentation throughout infected cultures, (**B**) variable degrees of fragmentation, (**C**) sarcomeres peeling away from nuclei, and (**D**) CMs missing nuclear DNA. All images are from 48 hours post infection with MOI = 0.006. White boxes indicate zoomed in areas. **E.** Comparison of intact CMs with SARS-CoV-2-infected CMs. On the right: schematic representation of the myofibrillar fragmetns observed, which retain an intact Z-disk and are broken along the contractile fibers. **F.** Imaging for mitochondrial integrity shows no overt immediate cellular apoptosis 72 hours post infection with low MOI (MOI = 0.006), but mitochondria are significantly herniated and appear apoptotic with high MOI (MOI = 0.1). **G.** Addition of the proteasome inhibitor bortezomib (1uM) induces similar myofibrillar breakdown as observed in SARS-CoV-2 infection, but less frequently and with diffuse cTnT signal. Left: representative immunofluorescence images (green: cTnT; blue: DAPI; red: dsRNA). Right: box plot with quantification of the number of cells presenting myofibrillar fragmentation (defined as at least one instance of an isolated cTnT doublet), normalized to total nuclei count. N=6. Each datapoint is the sum of nine randomly acquired fields of view. **: p-value<0.01; one-way ANOVA with Tukey's multiple comparisons.

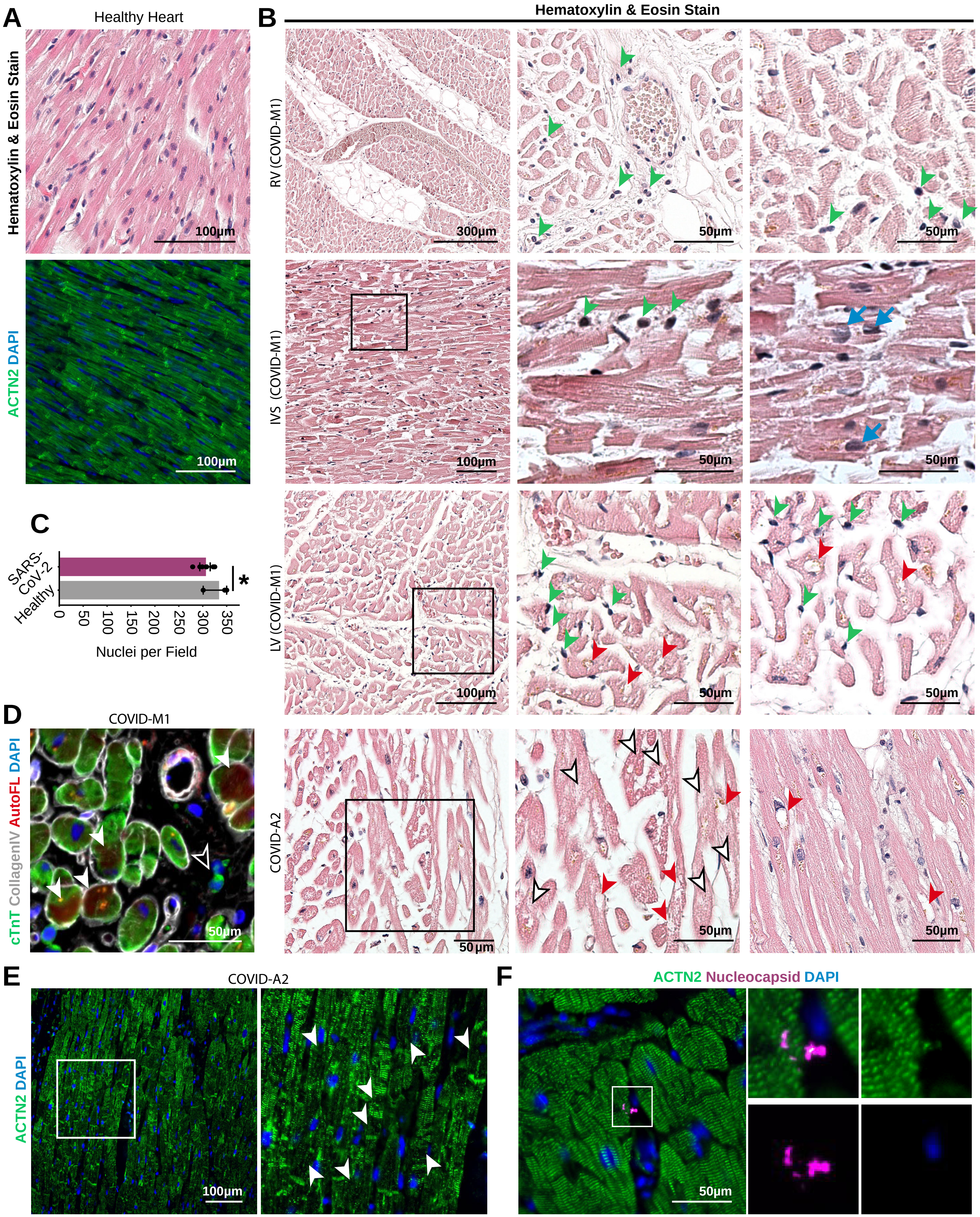

A
Healthy Heart
Hematoxylin & Eosin Stain
100µm
ACTN2 DAPI
100µm
B
Hematoxylin & Eosin Stain
RV (COVID-M1)
300µm
50µm
50µm
IVS (COVID-M1)
100µm
50µm
50µm
LV (COVID-M1)
100µm
50µm
50µm
COVID-A2
50 µm
50µm
50µm
C
SARS-CoV-2
Healthy
*
0
50
100
150
200
250
300
350
Nuclei per Field
D
COVID-M1
cTnT CollagenIV AutoFL DAPI
50µm
E
COVID-A2
ACTN2 DAPI
100µm
F
ACTN2 Nucleocapsid DAPI
50µm

**Supplementary Figure 6**. Additional representative images of COVID-19 patient hearts. **A.** Hematoxylin and eosin staining (H&E) (top) and immunohistochemistry (bottom) for ACTN2 in human healthy tissue evidencing densely packed cardiomyocyte myofibrils. **B.** H&E staining of right ventricle (RV), interventricular septum (IVS) and left ventricle (LV) from a COVID-19 patient with diagnosed myocarditis (COVID-M1) and of COVID-19 patients without diagnosed myocarditis (COVID-A2). Black boxes indicate zoomed inset areas of the figures at the right. White arrowheads indicate regions of sarcomeric disruption. Red arrowheads denote putative nuclear locations with loss of chromatin staining. Green arrowheads indicate mononuclear, potentially immune cells. Blue arrows denote enlarged and rounded cardiomyocyte nuclei. **C.** Quantification of nuclei per field of view of intact myocardium and disrupted myocardium from SARS-CoV-2 patients. *: p-val < 0.02.**D.** Left: Immunostaining of a COVID-19 patient heart with diagnosed myocarditis (COVID-M1), with irregular cardiac troponin T staining (white arrows) and the presence of small mononucleated cells in the interstitial tissue positive for cardiac troponin T (black arrows). **E.** Immunostaining of COVID-A2 patient hearts without diagnosed myocarditis display regions with extensive sarcomeric disruption, indicated with white arrowheads. **F.** Immunohistochemical staining for viral nucleocapsid protein (magenta) and α-actinin 2 (green) yielded no recognizable signal aside from occasional, unidentified puncta.
